# Tunable Extracellular Self-Assembly of Multi-Protein Conjugates from *Bacillus subtilis*

**DOI:** 10.1101/087593

**Authors:** Charlie Gilbert, Mark Howarth, Colin R. Harwood, Tom Ellis

## Abstract

The ability to stably and specifically conjugate recombinant proteins to one another is a powerful in vitro technique for engineering multifunctional enzymes, protein therapeutics and novel biological materials. However, for many applications spontaneous in vivo protein conjugation would be preferable to in vitro methods. Exploiting the recently described SpyTag-SpyCatcher system, we describe here how enzymes and structural proteins can be genetically-encoded to covalently conjugate in culture media following programmable secretion by Bacillus subtilis. Using this novel approach, we demonstrate how self-conjugation of a secreted industrial enzyme, XynA, dramatically increases its resilience to boiling and we show that cellular consortia can be engineered to self-assemble functional multi-protein complexes with tunable composition. This genetically-encoded modular system provides a new, flexible strategy for protein conjugation harnessing the substantial advantages of extracellular self-assembly.

---

In the context of biotechnology, proteins can be seen as modular components whose functions can be combined, augmented and refined by bringing them together to form complexes. Novel biological materials can be assembled^1,2^ and functionalised^3–5^ via linking proteins together and co-localizing enzymes from a single metabolic pathway can be used to enhance metabolic fluxes in biosynthesis^6,7^. In medical applications, vaccine efficacy can be improved by conjugating antigens to the surface of particles^8^ and therapeutic proteins can be stabilised or targeted to specific tissues and cells by fusing them to appropriate protein partners^9,10^. As a result, there is growing interest in methods to produce self-assembling protein-protein complexes for various applications.

Genetic fusion is a simple and direct method for conjugating proteins together, but is limited in both the size and topology of conjugates that can be formed^11^. Conversely, chemical conjugation methods enable multivalent and extensible protein-protein conjugation^12^, but typically require prior purification and treatment of proteins and so cannot be implemented *in vivo*. By contrast, biological conjugation methods such as the SpyTag-SpyCatcher system^13^, enable both genetically-programmed *in vivo* self-assembly and the formation of a variety of topologies^14,15^.

The SpyTag-SpyCatcher system^13^ directs specific, covalent conjugation of proteins through two short polypeptide tags: the SpyTag and SpyCatcher. The larger partner, the SpyCatcher, adopts an immunoglobulin-like fold which specifically binds the SpyTag and autocatalyses the formation of an intermolecular isopeptide bond between two amino acid side chains. Notably, in the few years since its initial description^13^ the SpyTag-SpyCatcher system has been applied to the production of programmable and customisable materials^4,16–18^, synthetic vaccines^8^, thermo-tolerant enzymes^19–21^, stably packaged enzymes^22,23^ and more^7,24,25^.

However, thus far, the SpyTag-SpyCatcher system has only ever been deployed within the cell or *in vitro*, following purification of individual components. Yet for a variety of applications, it would be advantageous for proteins to be secreted prior to conjugation. Such extracellular production greatly simplifies downstream processing and purification of products^26^, improving cost-effectiveness at industrial scale. In addition, secreting monomeric components of protein polymers avoids cytotoxicity commonly caused by their intracellular expression, facilitating applications such as protein material production^27^. Lastly, by engineering microbes to secrete proteins that complex together outside the cell, it becomes possible to compartmentalise the production of different proteins within different strains in a co-culture. Engineering so-called ‘cellular consortia’ to perform co-operative biological tasks in this way enables the division of labor between co-cultured strains, autonomous patterning of biomaterials and optimisation of biological processes^4,28–31^.

To provide a modular platform for these diverse applications, we sought to engineer simultaneous protein secretion and SpyTag-SpyCatcher-mediated protein conjugation using *Bacillus subtilis*, a Gram-positive bacterium used extensively in industrial biotechnology^32^ with considerable capacity for protein secretion (up to 20 grams per liter^33^). We designed and built recombinant fusion proteins consisting of separate protein modules specifying function, secretion and conjugation and demonstrate here that these modules are active and direct secretion and extracellular conjugation without perturbing enzymatic activity. Finally, we illustrate the utility of our system with two applications: producing secreted thermo-tolerant industrial enzymes and the spontaneous, tunable assembly of functional multi-protein complexes by cellular consortia.

## RESULTS AND DISCUSSION

### Expression, secretion and conjugation of SpyTag-SpyCatcher fusion proteins

To assess whether the SpyTag-SpyCatcher system could be coupled to *B. subtilis* protein secretion, we first fused together protein-encoding DNA modules specifying secretion, conjugation and function into single open reading frames (ORFs) within gene expression cassettes. Four protein modules, each connected by two amino acid glycine-serine linkers, were defined: an N-terminal secretion signal peptide, an upstream SpyPart – either SpyTag (T) or SpyCatcher (C) – then a user-defined protein of interest and a C-terminal His_6_-tagged SpyPart (Figure 1A).

**Figure 1.**
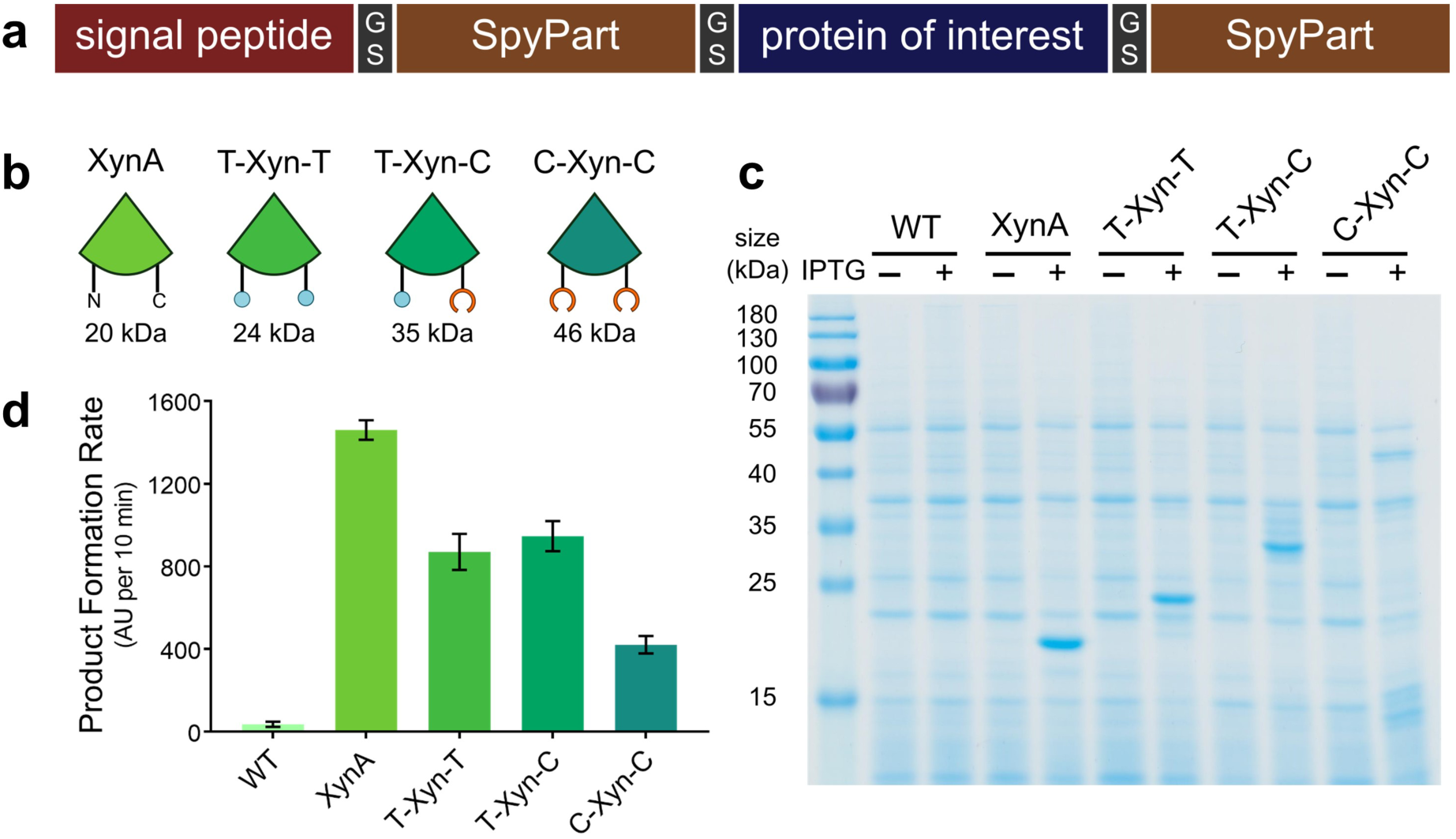
Design, secretion and activity of SpyTag-SpyCatcher XynA fusion proteins. (**a**) Protein-encoding DNA modules specifying secretion, conjugation and function were fused into single ORFs separated by 2-amino acid glycine-serine linkers. (**b**) Four recombinant proteins based on XynA were designed: the full-length xylanase XynA, SpyTag-XynA-SpyTag (T-Xyn-T), SpyTag-XynA-SpyCatcher (T-Xyn-C) and SpyCatcher-XynA-SpyCatcher (C-Xyn-C). All possess the native XynA signal peptide at their N-termini as well as a C-terminal His_6_ tag. (**c**) Culture supernatants from uninduced (-) and IPTG-induced (+) cultures were 10x concentrated by TCA precipitation prior to SDS-PAGE analysis with Coomassie staining. All four proteins were well-expressed and secreted. Untransformed *B. subtilis* WB800N (WT) was included as a negative control. (**d**) Culture supernatants were analysed for xylanase activity using a fluorogenic substrate. All samples exhibited xylanase activity, indicating secretion of active fusion proteins. Product formation rates were calculated over 10 minutes from triplicate samples (data represent the mean ± 1 SD).

Using this design, we first generated a series of fusion proteins based on the native, secreted *B. subtilis* endo-xylanase, XynA (Figure 1B and Supplementary Figure 1), a hemicellulose-degrading enzyme with uses in industry. The native XynA signal peptide was preserved at the N-terminus and SpyParts were fused either side of the XynA enzyme core to create three proteins: T-Xyn-T, T-Xyn-C and C-Xyn-C (Figure 1B). As a control, a construct expressing the full-length XynA with a C-terminal His_6_-tag was also created. All constructs were cloned downstream of the strong IPTG-inducible P_grac_ promoter in pHT01, a *B. subtilis-E. coli* shuttle vector. Each of these fusion proteins was successfully expressed and secreted from *B. subtilis* (Figure 1C and Supplementary Figure 2) and retained xylanase activity (Figure 1D).

To verify the activity of the secreted SpyTag and SpyCatcher motifs, we first purified the His_6_-tagged T-Xyn-T and C-Xyn-C proteins from *B. subtilis* culture supernatant using immobilised metal ion affinity chromatography (IMAC) (Supplementary Figure 3). Purified proteins were mixed (Figure 2A) and analysed by Western blot (Figure 2B). Immediately upon mixing, covalently-conjugated polymeric species were formed (Figure 2B) indicating that SpyTag and SpyCatcher were functional. After longer periods of incubation, the majority of monomers were converted to a polymeric form (Supplementary Figure 4).

**Figure 2.**
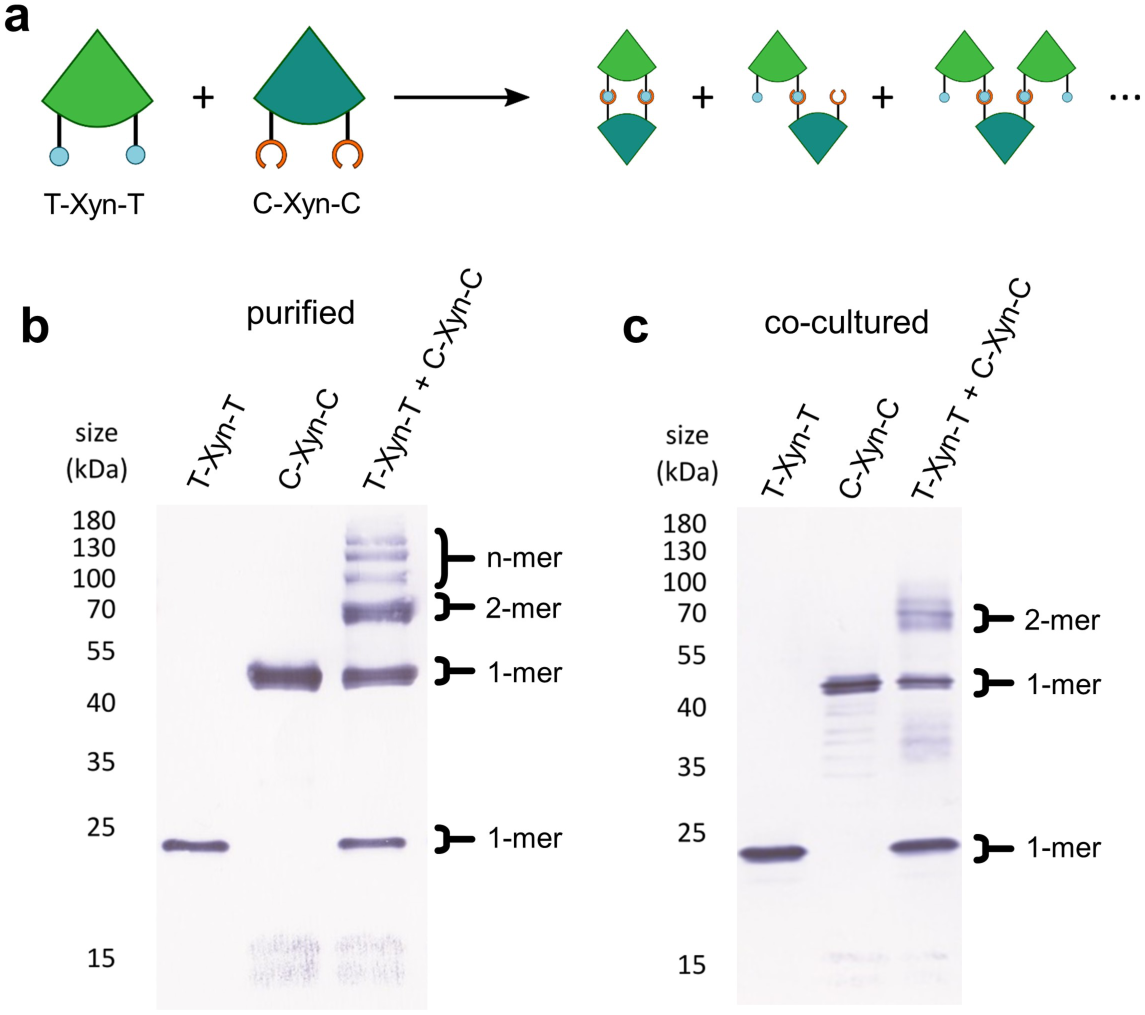
SpyTag-SpyCatcher-mediated protein conjugation from co-cultures. (**a**) Schematic showing the possible reaction products between T-Xyn-T and C-Xyn-C. (**b**) His_6_-tagged T-Xyn-T and C-Xyn-C fusion proteins were purified from culture supernatants by IMAC (immobilised metal ion affinity chromatography) and analysed by Western blotting (with an anti-His_6_ antibody) in isolation or after mixing. (**c**) Supernatant samples from monocultures expressing T-Xyn-T or C-Xyn-C and a co-culture of T-Xyn-T and C-Xyn-C were analysed by Western blotting with an anti-His_6_ antibody. A dimeric species was clearly discernible under co-culture conditions, along with monomers of T-Xyn-T and C-Xyn-C.

To determine whether SpyTag and SpyCatcher were active under co-culture, strains expressing T-Xyn-T and C-Xyn-C were grown alone or together and supernatant samples analysed by Western blot (Figure 2C). A species with mobility corresponding to a dimer was detected after two hours of co-culture (Figure 2C), indicating that SpyTag and SpyCatcher are indeed functional under co-culture conditions. Detection of polymeric species over longer periods of co-culture was visible, but hampered by inherent proteolysis from the two extracellular proteases native to *B. subtilis* WB800N and by smearing of bands at higher molecular weights (Supplementary Figure 5).

To verify that the secreted T-Xyn-T and C-Xyn-C proteins were able to conjugate under co-culture, we purified His_6_-tagged species from the supernatant of a co-culture of strains expressing both T-Xyn-T and C-Xyn-C and their approximate molecular weights determined through size exclusion chromatography-multi-angle light scattering (SEC-MALS) analysis (Supplementary Figure 6). Consistent with our previous observations, the major species present exhibited a molecular weight corresponding to a T-Xyn-T and C-Xyn-C dimer. In addition, oligomer species were also detected.

### Engineering XynA thermo-tolerance by SpyRing cyclisation

To demonstrate the utility of this technique for protein engineering, we tested the ability of the SpyTag-SpyCatcher reaction to improve the thermo-tolerance of XynA through SpyRing cyclisation. Protein cyclisation by linkage of the N- and C-termini has been shown to increase the ability of enzymes to tolerate exposure to high temperatures – are highly-desirable trait for many industrial enzymes – and can been achieved through a number of methods^34–36^. The SpyRing system works through fusion of the SpyTag and SpyCatcher respectively at the N- and C-termini of a protein, leading to covalent cyclisation that dramatically improves the ability of globular proteins to refold to native structures following exposure to high temperatures^37^.

In addition to strains secreting the full length XynA and T-Xyn-C proteins, we engineered a strain to secrete a protein bearing the mutated SpyCatcher^E77Q^ (C’) unable to form the covalent isopeptide linkage with the SpyTag^13^, T-Xyn-C’ (Figure 3A-C). Due to the close proximity of the N- and C-termini of XynA (within 1 nm, Supplementary Figure 1) we anticipated that the SpyTag and SpyCatcher of the T-Xyn-C protein would be capable of reacting intramolecularly to cyclise XynA. While competing polymerisation reactions are also possible (Figure 3C), we expected cyclisation to be the major product, as seen with other SpyRing cyclisations^19^, particularly since the concentration of the T-Xyn-C protein is relatively low in the culture medium.

**Figure 3.**
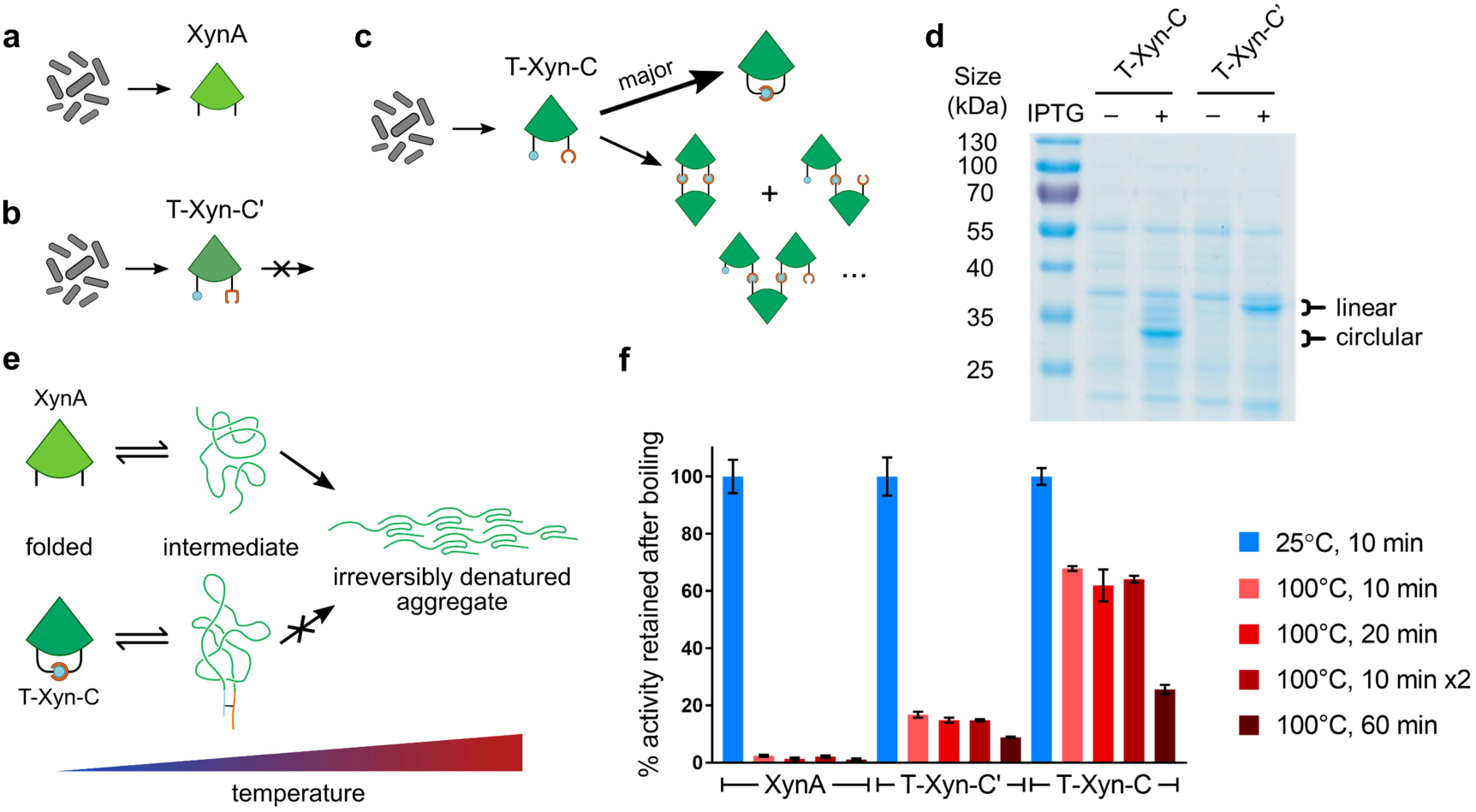
SpyRing cyclisation confers XynA thermo-tolerance. Strains expressing XynA (**a**), the mutant SpyTag-XynA-SpyCatcher^E77Q^ (T-Xyn-C’) (**b**) and SpyTag-XynA-SpyCatcher (T-Xyn-C) (**c**) were created. The T-Xyn-C protein is able to cyclise through SpyRing cyclisation. (**d**) Comparison of the electrophoretic mobility of T-Xyn-C and T-Xyn-C’ proteins by SDS-PAGE with Coomassie staining (expected molecular mass ∼ 35 kDa). A clear difference in electrophoretic mobility of the two proteins is seen, consistent with covalent cyclisation of T-Xyn-C. (**e**) SpyRing cyclisation works by covalently conjugating the N and C-termini of a globular protein. Upon boiling, the folded protein begins to unfold. The SpyRing system is believed to prevent the irreversible transition of partially unfolded intermediates to denatured aggregates and therefore enable refolding when the temperature is lowered. (**f**) Culture supernatants from strains expressing XynA, T-Xyn-C’ and T-Xyn-C were subjected to a variety of high-temperature programs and subsequently assayed for xylanase activity. The T-Xyn-C protein shows dramatically increased tolerance to a variety of high temperature programs compared to XynA and T-Xyn-C’. Samples were analysed in triplicate, data represent the mean ± 1 SD.

To confirm cyclisation, which is known to perturb the mobility of proteins during gel electrophoresis^19^, we compared the electrophoretic mobility of T-Xyn-C to that of the mutant T-Xyn-C’. Consistent with SpyRing cyclisation, the T-Xyn-C and T-Xyn-C’ proteins exhibited substantially different mobilities under gel electrophoresis (Figure 3D and Supplementary Figure 3). To next determine whether SpyRing cyclisation confers thermo-tolerance to XynA by preventing irreversible aggregation (Figure 3E) we subjected supernatant samples from cultures secreting the native XynA, T-Xyn-C’ and T-Xyn-C proteins to a variety of high-temperature conditions. After cooling to 4°C these samples were then assayed for xylanase activity (Figure 3F). All supernatants exhibited similar levels of xylanase activity following incubation at 25°C (Supplementary Figure 7). However, following exposure to high-temperature conditions only supernatants containing T-Xyn-C retained substantial levels of xylanase activity (Figure 3F). Remarkably after exposure to 100°C for 10 min, T-Xyn-C retained 67.9% ±0.9 of its xylanase activity in contrast to negligible activity (2.4% ±0.3) for XynA, and a similar protective effect was also seen across a variety of other high-temperature programs. Consistent with previous studies^19^, a mild protective effect was also observed for the mutant control T-Xyn-C’. This is likely due to the relatively strong, non-covalent interactions between SpyTag and SpyCatcher mutants^13^.

Owing to its ease of implementation and the availability of guidelines for its design^40^, the SpyRing system is an attractive tool for improving the stability of enzymes. The SpyRing cyclisation system has previously been harnessed to improve the thermo-tolerance of a number of other intracellularly expressed enzymes with industrial relevance^20,21^. However, since extracellular production of proteins for biotechnology vastly improves cost-effectiveness, our strategy offers a novel, attractive approach to the production and stabilisation of industrially-relevant enzymes. Notably, xylanases with improved thermal stability are of great interest to industry, offering an eco-friendly alternative to the chemicals used in the paper pulp bleaching process^42^ and as an additive to improve animal feed digestibility^43^. And further, *B. subtilis* naturally secretes a number of other enzymes with uses industry, including amylases, proteases and lipases.

### A modular method for multi-protein complex assembly from cellular consortia

Having demonstrated secretion and conjugation of XynA fusion proteins, we next looked to exploit this approach to create extracellular multi-protein complexes from engineered cellular consortia. To facilitate assembly of plasmid constructs, we first designed a Golden Gate assembly system (Supplementary Figure 8A). This strategy allowed simple, one-step assembly of ORFs encoding an N-terminal signal peptide for secretion, an upstream SpyPart, a user-defined protein of interest and a C-terminal His_6_-tagged SpyPart. Parts were initially cloned into entry vectors from which they can be verified, stocked and re-used for future assemblies (Supplementary Figure 8B). Stocked parts were then assembled directly into the pHT01 IPTG-inducible expression vector ready for use in *B. subtilis*.

Using this system, we created a series of plasmid constructs for secretion of recombinant proteins based on a second native *B. subtilis* enzyme, the endo-cellulase, CelA (Supplementary Figure 9). Like xylanases, cellulases have attracted interest in variety of industrial contexts, notably for their ability to degrade plant biomass, a sustainable potential feedstock for bio-commodity production^38,39^. In fact, creating multi-enzyme complexes of synergistic plant biomass-degrading enzymes such as CelA and XynA, has previously been shown to enhance the degradation of complex cellulosic substrates^40^. As with XynA, we found that SpyParts could be fused to the N- and C-termini of CelA without disrupting secretion (Supplementary Figure 9B and 9C) or enzyme activity (Supplementary Figure 9D). Further, these SpyParts were active and covalently conjugated XynA and CelA fusion proteins under co-culture to form multi-enzyme complexes (Supplementary Figure 10).

To demonstrate the compatibility of heterologous proteins with our system, we created a plasmid construct for expression and secretion of an elastin-like polypeptide (ELP) fused to SpyParts. The ELP used here, ELP_20-24_, is a short 10 kDa polypeptide derived from human tropoelastin, consisting of one hydrophilic domain flanked by two hydrophobic domains^41^. Owing to its short size, ELP_20-24_, does not undergo coacervation under conditions used here.

A tagged ELP_20-24_ protein, T-ELP-T, was generated with an N-terminal signal peptide from the *B. subtilis* SacB protein, an upstream SpyTag, a downstream SpyTag and a C-terminal His_6_ tag (Supplementary Figure 11A and Figure 4A). A second version of T-ELP-T was generated in which the isopeptide bond-forming aspartate residue of each SpyTag was mutated to alanine (T’-ELP-T’-H), preventing covalent conjugation with SpyCatcher^13^ (Supplementary Figure 11A and Figure 4A). In addition, plasmids expressing the C-Xyn-C and C-Cel-C proteins were also modified to remove their C-terminal His_6_ tags.

Strains expressing T-ELP-T and the mutant T’-ELP-T’ were then co-cultured with strains expressing the C-Xyn-C and C-Cel-C. Supernatant samples from monocultures of each of the strains and from three-strain co-cultures were then analysed by Western blot with an anti-His_6_ antibody (Supplementary Figure 11B). The T-ELP-T and the mutant T’-ELP-T’ proteins were well-expressed and secreted. When cultured together with the C-Xyn-C and C-Cel-C proteins, the mobility of the mutant T’-ELP-T’ was unaffected, whereas almost all of the secreted T-ELP-T protein was incorporated into dimeric and polymeric species (Supplementary Figure 11B), verifying conjugation by the Spy system.

The ability to co-localise cooperative enzymes – ones that act in concert on a single substrate or in a pathway – has been shown to improve metabolic fluxes and is consequently a useful approach in metabolic engineering^6,7^. Indeed, this is an strategy employed in nature; certain bacteria able to metabolise plant biomass produce large, extracellular multi-protein complexes known as cellulosomes, consisting of numerous synergistically-acting enzymes^44^. Notably, in efforts to engineer recombinant microbes capable of growth on plant biomass, two- and three-protein ‘designer-cellulosomes’ have previously been assembled *in vitro* and by co-culturing protein-secreting bacterial strains^45,46^. Although in contrast to our system, these complexes were assembled through the cohesin-dockerin interaction, a non-covalent protein-protein interaction^47^.

**Figure 4.**
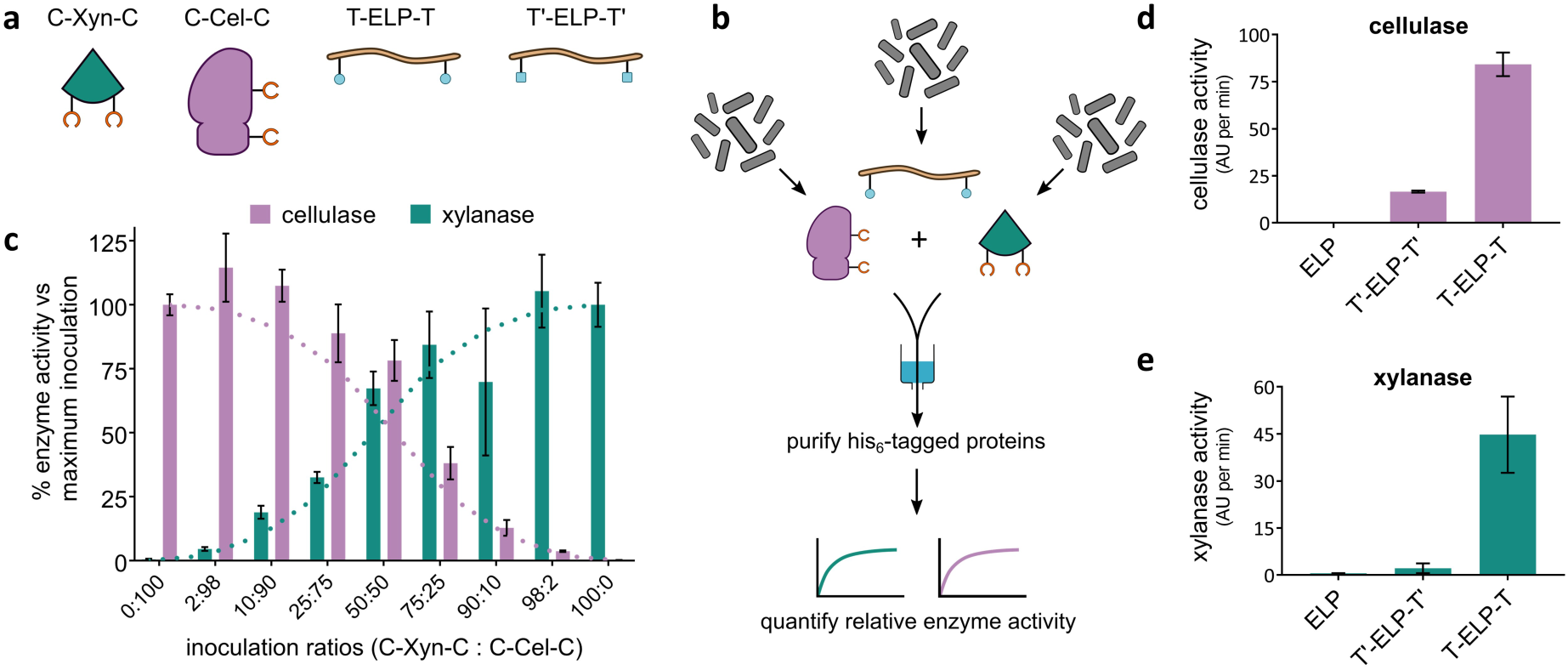
Tuning the composition of a multi-protein complex. (**a**) The four recombinant proteins used here: SpyCatcher-XynA-SpyCatcher lacking a C-terminal His_6_ tag (C-Xyn-C), SpyCatcher-CelA-SpyCatcher lacking a C-terminal His_6_ tag (C-Cel-C), SpyTag-ELP_20-24_-SpyTag-His_6_ (T-ELP-T) and SpyTag^DA^-ELP_20-24_-SpyTag^DA^-His_6_ (T’-ELP-T’). Molecular weights of each species are also given (calculated assuming removal of N-terminal signal peptides). (**b**) Schematic illustrating the co-purification experiment. (**c**) By modulating the proportion of cells expressing the C-Xyn-C (teal dotted line) inoculated compared to the proportion of cells expressing the C-Cel-C (lilac dotted line) inoculated, the proportions of cellulase and xylanase activities co-purified with T-ELP-T could be tuned across a variety of different inoculation ratios (samples prepared in triplicate, data represent the mean ± 1 SD). Co-purification of cellulase (**d**) and xylanase (**e**) activity was repeatable and dependent on the SpyTag-SpyCatcher reaction, as co-purification sharply decreased when performed with an ELP construct lacking SpyTags (ELP) or with mutated SpyTags (T’-ELP-T’). Samples prepared in triplicate, data represent the mean ± 1 SD.

### Tuning multi-protein complexes using consortia composition

One of the great advantages of engineering cellular consortia to carry out a biological process, is that the balance between different sub-processes can be tuned simply by tuning the relative productivity of different strains within the co-culture. We therefore set out to tune the relative amounts of C-Xyn-C and C-Cel-C incorporated into multi-protein complexes with T-ELP-T, simply by tuning their relative inoculation ratios in three-strain co-cultures. To quantify the relative amounts of C-Xyn-C and C-Cel-C in multi-protein complexes, we performed co-purifications using the C-terminal His_6_ tag fused to ELP proteins. As outlined in Figure 4B, following three-strain co-culture growth, His_6_-tagged ELP proteins were isolated from the culture supernatant by IMAC purification, and any SpyTag-SpyCatcher conjugated proteins were co-purified along with them while unbound proteins were washed off. The relative levels of C-Xyn-C and C-Cel-C incorporated into multi-protein complexes were then quantified via enzyme activity assays.

We performed several three-strain co-cultures in which the inoculum volume of the T-ELP-T expressing strain was fixed and the inoculation proportions of C-Xyn-C and C-Cel-C expressing strains varied. Enzyme activity was detected in all purified fractions, demonstrating the formation of functional multi-protein complexes. Remarkably, we observed that the proportions of CelA and XynA proteins incorporated into the extracellular multi-protein conjugates could be finely tuned simply by adjusting the proportions of the strains in the initial inoculations (Figure 4C). Furthermore, the relative enzyme activities of these complexes matched the relative inoculation proportions closely over a range of conditions. Additional co-purifications with two negative control strains: a strain expressing the mutant T’-ELP-T’ only capable of non-covalent binding and a strain expressing secreted ELP-H lacking SpyTags, verified that the conjugation between all three proteins in the culture medium was specifically SpyTag-SpyCatcher-mediated as both controls showed dramatically reduced cellulase (Figure 4D) and xylanase (Figure 4E) levels. We thus verified our ability to tune the relative proportions of XynA and CelA incorporated into multi-protein complexes simply by tuning their relative inoculation ratios.

This system thus offers a simple way to both assemble functional multi-protein complexes and to fine-tune their properties. This approach could be useful in any scenario in which the proportions of individual components of multiprotein complexes influences the desired functions. For instance, when co-localising co-operative enzymes to improve flux through a metabolic pathway – as with the previously-mentioned ‘designer cellulosomes’ – tuning enzyme proportions may enable improved yields by increasing the levels of enzymes catalysing rate-limiting steps or decreasing the levels of enzymes producing toxic pathway intermediates.

## CONCLUSION

Here we have demonstrated the feasibility and utility of combining protein secretion with SpyTag-SpyCatcher-mediated protein conjugation. As an initial illustration of the utility of our method, we coupled SpyRing cyclisation with protein secretion, enabling one-step extracellular production and stabilisation of the endo-xylanase, XynA. Additionally, we applied our method to engineer extracellular production of self-assembling multi-protein complexes from cellular consortia. Our approach allows the relative proportions of proteins incorporated into multi-protein complexes to be tuned simply by varying their relative inoculation ratios in co-cultures, rather than requiring any additional genetic engineering such as promoter swapping.

Beyond the work presented here, the productivity of different strains within co-cultures could be further controlled by coupling expression with additional genetic circuits, such as inducible switches and quorum sensing systems^48^. Indeed, these tools have been previously harnessed within cellular consortia to program temporal and spatial control over monomer patterning within amyloid fibrils^4^. In addition, transferring the strategy to alternative secretion hosts – such as the yeasts *Saccharomyces cerevisiae* and *Pichia pastoris* – would enable the secretion of a much broader range of heterologous proteins, relieving the need for compatibility with *B. subtilis*. As our approach is modular in design there is also great scope for integrating further components to broaden potential applications, such as alternative functional components or biological protein conjugation methods^15,47,49,50^. Lastly, while the work here focuses exclusively on bivalent proteins – those possessing two SpyParts – incorporating additional SpyParts into fusion proteins can enable the formation of extended, branching polymeric networks and hydrogels^16,18^.

Programming protein conjugation and self-assembly within the extracellular environment offers great promise in the effort to generate novel industrial enzymes, multi-protein complexes and biological materials: improving production cost-effectiveness, reducing cellular burden and toxicity and enabling patterning and tunability through engineered cellular consortia. The modular approach described here therefore offers a platform for the development of biotechnological products to meet real-world challenges.

## Acknowledgements

We are grateful to Dr Carlos Bricio-Garberi and Dr Olivier Borkowski for constant advice, discussions and suggestions. We thank Christopher Sauer and Rita Cruz for providing advice and guidance on *B. subtilis* biology and protein secretion, Dr Christopher Schoene for advice regarding SpyRing cyclisation, Dr Alex Webb for providing protein expression strains and constructs and Dr Ciaran McKeown for performing the SEC-MALS experiment.

## METHODS

### Strains and Plasmids

Bacterial plasmids and strains used in this study are listed in Supplementary Table 1 and Supplementary Table 2, respectively. Both *B. subtilis* and *E. coli* were grown in LB medium or 2xYT medium at 37°C under aeration. In all instances media were supplemented with appropriate antibiotics at the following concentrations for *E. coli*: ampicillin 100 µg.ml^−1^, chloramphenicol 34 µg.ml^−1^, kanamycin 50 µg.ml^−1^. For *B. subtilis*, media were supplemented with 5 µg.ml^−1^ chloramphenicol.

### lasmid construction

All plasmids constructed in this study were constructed using standard cloning techniques. Oligonucleotides were obtained from IDT. Restriction endonucleases, Phusion-HF DNA polymerase and T7 DNA ligase were obtained from NEB. Unless stated, all plasmids were transformed into *E. coli* turbo (NEB) for amplification and verification before transforming into *B. subtilis* WB800N for protein expression and secretion. All constructs were verified by restriction enzyme digestion and Sanger sequencing (Source Bioscience). Amino acid sequences of protein parts used in this study are given in Supplementary Table 3.

To create the pHT01-xynA-His_6_ construct, the native *B. subtilis* xynA ORF was amplified from the genome of *B. subtilis*168 by colony PCR. Oligonucleotides were designed to introduce a C-terminal His_6_ tag as well as upstream *BamHI* and downstream *AatII* restriction enzyme sites. The amplified xynA-His_6_ ORF and pHT01 backbone were digested with BamHI and AatII and gel purified and T4-ligated.

Using pHT01-xynA-His6 as a starting point, Golden Gate assembly was used to construct pHT01-xynA_SP_-SpyTag-xynA-SpyTag-His_6_ (T-Xyn-T), pHT01-xynA_SP_-SpyTag-xynA-SpyCatcher-His_6_ (T-Xyn-C) and pHT01-xynA_SP_-SpyCatcher-xynA-SpyCatcher-His_6_ (C-Xyn-C). Two versions of the SpyCatcher were synthesised by GeneArt (Life Technologies). A set of SpyCatcher-coding sequences codon-optimised for *B. subtilis* were created and the two most divergent sequences chosen (this was to reduce the risk of recombination within constructs containing two copies of the SpyCatcher). Two versions of the SpyTag were codon-optimised in the same manner and created from overlapping oligonucleotides. Golden Gate assemblies of gel purified PCR products using BsaI were performed as described^51^. SpyTag and/or SpyCatcher sequences were introduced between the xynA signal peptide (xynA_SP_) and xynA enzyme and between the xynA enzyme and His_6_ tag. In each instance, 4 bp overhangs were incorporated into glycine-serine (GS) linkers. The backbone was amplified in two halves to allow mutation (and therefore removal) of an unwanted BsaI site in the AmpR cassette.

The pHT01-xynA_SP_-SpyTag-xynA-SpyCatcher^E77Q^-His_6_ (T-Xyn-C’) mutant construct was created using pHT01-xynA_SP_-SpyTag-xynA-SpyCatcher-His_6_ (T-Xyn-C) as a template. We used BsaI Golden Gate assembly-based mutagenesis to mutate the catalytic glutamate of SpyCatcher to glutamine.

To suit our cloning needs we created a modular DNA assembly toolkit based on Golden Gate assembly (Supplementary Figure 8A). Four separate ORF parts were defined: a signal peptide part, an upstream SpyPart, a central protein of interest part and a downstream SpyPart. Each position was defined by the sequence of specific 4 bp overhangs generated by BsaI digestion upstream and downstream of the part. Where fewer than four ORF parts are desired in the final construct, the 4 bp overhangs can be modified accordingly. ORF parts were cloned into a Golden Gate assembly part vector, pYTK001, where they were sequence-verified and stocked for subsequent assemblies. Stocked parts used in this study are summarised in Supplementary Figure 8B, their sequences given in Supplementary Table 4 and are available from Addgene.

We also created an entry vector derived from pHT01, called pCG004, itself assembled by BsaI Golden Gate assembly. The pHT01 backbone was amplified by PCR – again in two halves to allow removal of the unwanted BsaI site – and a dropout part introduced downstream of the P_grac_ promoter and upstream of the terminator. The dropout part consists of a constitutive GFP mut3b^52^ expression cassette flanked by BsaI restriction sites. Successful Golden Gate assembly will result in removal of the GFP expression cassette and therefore visual (green-white) screening of transformants. The GFP expression cassette was created using the P_veg_ promoter and spoVG RBS, specifically chosen for their activity in both *E. coli* and *B. subtilis* – and therefore allowing transformation of Golden Gate assemblies into either strain.

We used our Golden Gate assembly system to construct pHT01-celA-his_6_ (CelA), pHT01-celA_SP_-SpyTag-celA-SpyTag-His_6_ (T-Cel-T), pHT01-celA_SP_-SpyTag-celA-SpyCatcher-His_6_ (T-Cel-C) and pHT01-celA_SP_-SpyCatcher-celA-SpyCatcher-His_6_ (C-Cel-C). To minimise the size of these constructs we used SpyCatcherΔN1ΔC2, which has superfluous amino acids trimmed from its N- and C-termini^53^ (mSpyCatcher). Since repeated attempts to clone the pHT01-celA_SP_-SpyCatcher-celA-SpyCatcher-His_6_ (C-Cel-C) plasmid into *E. coli* resulted in identical mutations of the upstream SpyCatcher, it was cloned directly into *B. subtilis* WB800N and sequence-verified. We also used our Golden Gate assembly system to construct pHT01-sacB_SP_-ELP_20-24_-His_6_ (ELP) and pHT01-sacB_SP_-SpyTag-ELP_20-24_-SpyTag-His_6_ (T-ELP-T).

BsaI Golden Gate assembly-based mutagenesis was used to construct: the mutated pHT01-sacB_SP_-SpyTag^DA^-ELP_20-24_-SpyTag^DA^-His_6_ (T’-ELP-T’) (from pHT01-sacB_SP_-SpyTag-ELP_20-24_-SpyTag-His_6_), the His_6_ tag-lacking pHT01-xynA_SP_-SpyCatcher-xynA-SpyCatcher (from pHT01-xynA_SP_-SpyCatcher-xynA-SpyCatcher-His_6_) and the His_6_ tag-lacking pHT01-celA_SP_-SpyCatcher-celA-SpyCatcher (from pHT01-celA_SP_-SpyCatcher-celA-SpyCatcher-His_6_). Similar to the construct from which it was derived, pHT01-celA_SP_-SpyCatcher-celA-SpyCatcher repeatedly showed mutations when cloned into *E. coli* and so was instead cloned directly into *B. subtilis* WB800N and sequence-verified.

### Protein expression and co-culturing

In all instances, glycerol stocks of *Bacillus subtilis* strains were first spread onto selective LB plates from which single colonies were used to inoculate 5 mL 2xYT liquid cultures. After 16 h of growth, strains were back-diluted 1/50 into 5 mL of fresh 2xYT medium. Where indicated, protein expression was induced with 1 mM IPTG. Expression culturing was performed for between 2 h and 8 h, depending on the individual experiment. To collect secreted protein fractions, cultures were centrifuged at 3220 x g for 10 min and supernatants harvested.

### SDS-PAGE and Western blotting

Since the concentration of proteins in the culture supernatant is relatively low, trichloroacetic acid (TCA) precipitation was performed to concentrate samples (by a factor of 10) prior to analysis by SDS-PAGE and Western blotting. Secreted proteins in the supernatant were precipitated by adding 100 µL of 4°C 100% TCA to 900 µL of culture supernatant and incubating for 16 h at 4°C. Precipitated proteins were centrifuged at 16900 x g for 10 min at 4°C, washed with 1 mL of ice-cold acetone, centrifuged again at 16900 x g for 10 min at 4°C and finally air-dried. Protein-containing pellets were then resuspended in 90 µL of 1x SDS-PAGE sample buffer (0.2 M Tris-HCl pH 6.8, 2% SDS, 10% glycerol, 0.01% bromophenol blue + 2 mM DTT) and boiled for 10 min.

SDS-PAGE gels – with differing separating gel percentages depending on the size of proteins analysed – were run as standard and proteins stained using SimplyBlue SafeStain (Thermo). Alternatively, proteins were transferred to a PVDF membrane for immunodetection using a mouse anti-His_6_ primary antibody (BioLegend clone: J099B12) and an alkaline phosphatase-conjugated anti-mouse secondary antibody (Promega). Bound antibodies were detecting using a BCIP-NBT colorimetric kit (Life Technologies).

### Enzyme activity assays

Assays for xylanase and cellulase activities were performed using the EnzChek Cellulase Substrate (Thermo) and EnzChek Ultra Xylanase Assay Kit (ThermoFisher). Substrate solutions were prepared according to the manufacturer’s instructions. In both assays 50 μL supernatant samples were pipetted into a Costar 96-well Cell Culture Plate (Corning) and 50 μL substrate solutions added simultaneously with a multi-channel pipette. Samples were immediately analysed on a Synergy HT plate reader – both assays report enzyme activity through a fluorogenic substrate. All assays were performed at room temperature with biological triplicates. Data are represented as plots of showing the accumulation of fluorescent product over time or by calculating the enzyme reaction rate (gradient of the linear region of the graph).

### Protein purification

Protein purifications were performed using HisPur Ni-NTA Spin Columns (ThermoFisher), with 0.2 mL resin bed volume, according to the manufacturer’s instructions. Prior to purification, 3 mL samples of culture supernatant were first mixed with a 10x concentrated equilibration solution (500 mM NaH_2_PO_4_, 2.15 M Sodium Chloride, 100 mM imidazole, pH 8.0) to enable efficient binding of His6-tagged proteins to the Ni-NTA resin. A total of 2 mL of supernatant was passed over the Ni-NTA resin in three batches of 666 μL, with each incubated with the resin for 15 min prior to collecting flow through. Three washes were performed with 666 μL of wash buffer (50 mM NaH_2_PO_4_, 300 mM Sodium Chloride, 20 mM imidazole, pH 8.0) followed by elution in 500 μL of elution buffer (50 mM NaH_2_PO_4_, 300 mM Sodium Chloride, 250 mM imidazole, pH 8.0).

### SEC-MALS sample preparation

To prepare more concentrated purified protein required for SEC-MALS analysis, 5 mL seed cultures of strains expressing T-Xyn-T and C-Xyn-C were inoculated into 200 mL 2xYT medium to a final OD_600_ = 0.1 and incubated at 37 °C. When an OD_600_ ∼0.5 was reached, protein expression was induced by the addition of IPTG to a final concentration of 1 mM. After a further 4 h incubation, the culture was centrifuged at 4000 x g at 4° C for 20 min. The supernatant was removed and again centrifuged at 4000 x g at 4° C for 20 min. 150 mL of clarified supernatant was harvested. To this, 16.7 mL of 10x equilibration buffer (500 mM NaH_2_PO_4_, 2.15 M Sodium Chloride, 100 mM imidazole, pH8.0) was added. 10 mL of pre-equilibrated HisPur Ni-NTA resin (Thermo) was then added to the supernatant and incubated on ice with shaking for 1 h. The resin was allowed to settle out on ice and ∼130 mL of supernatant removed by pipetting. The resin was then resuspended with the remaining liquid and transferred to a 10 mL Pierce Disposable Column (Thermo) and the supernatant eluted off the column. Three washes were performed with 10 mL of wash buffer (50 mM NaH_2_PO_4_, 300 mM Sodium Chloride, 20 mM imidazole, pH8.0) followed by two 10 mL elutions with elution buffer (50 mM NaH_2_PO_4_, 300 mM Sodium Chloride, 250 mM imidazole, pH8.0). Elution fractions were concentrated to ∼2 mg.ml^−1^ using Pierce Protein Concentrator columns (10K MWCO, Thermo).

### Enzyme thermo-tolerance assays

To assess the ability of different proteins to withstand boiling, supernatants samples were exposed to specified temperature programs using a ProFlex PCR System (ThermoFisher) thermal cycler. Following boiling, samples were cooled a rate of 3 °C/sec to 4°C, re-equilibrated to room temperature and assayed for xylanase activity. Data were presented as plots showing the accumulation of fluorescent product over time. In addition, reactions rates were calculated by taking gradient over the linear (early) region of the fluorescence-time plots. The percentage of enzyme activity retained after boiling was calculated by comparing reactions rates between samples subjected to 25 °C for 10 min and samples subjected to 100 °C for 10 min (or otherwise stated).

### Co-culture conditions, co-purifications and enzyme assays

Two-strain co-cultures were performed with strains expressing SpyTag-CelA-SpyTag-His_6_ and SpyCatcher-XynA-SpyCatcher proteins. Seed cultures in 2xYT medium were grown for 16 h and used to inoculate co-cultures. Inductions were performed by inoculating 5 mL 2xYT medium containing 1 mM IPTG with 50 μL of each strain, or 100 μL of each strain for monocultures. Supernatant samples were harvested after 6 h and 8 h of incubation at 37 °C and analysed by SDS-PAGE and Western blotting.

Three-strain co-cultures were performed with strains expressing C-Xyn-C, C-Cel-C and various ELP_20-24_-containing proteins. In this instance triplicate seed cultures in 2xYT medium were grown for 16 h. Since seed cultures consistently reached similar optical densities (OD_600_ = 4.5 − 5.5), identical inoculation volumes were used for each replicate during inductions.

To compare the ability of the T-ELP-T, the mutant T’-ELP-T’ and the ELP proteins to co-purify xylanase and cellulase activities, 40 μL of each of the three strains were inoculated into 5 mL 2xYT medium containing 1 mM IPTG. After 8 h of incubation at 37 °C supernatants were harvested and analysed by Western blotting using an anti-His_6_ antibody. In addition, IMAC purifications of His_6_-tagged proteins in the supernatant were performed as described above.

To test the ability to tune the composition of multi-protein complexes, a number of different three-strain co-cultures were prepared in which the ratio of strains expressing the C-Xyn-C and C-Cel-C proteins was varied. The T-ELP-T-expressing strain inoculation was fixed at 40 μL. The total inoculation volume of strains expressing the C-Xyn-C and C-Cel-C proteins was fixed at 80 μL and varied: 0%:100% (0 μL:80μL), 2%:98% (1.6 μL:78.4 μL), 10%:90% (8 μL:72 μL), 25%:75% (20 μL:60 μL), 50%:50% (40 μL:40 μL) 75%:25% (60 μL:20 μL), 90%:10% (72 μL:8 μL), 98%:2% (78.4 μL:1.6 μL) and 100%:0% (80 μL:0 μL). After 8 h of incubation at 37 °C supernatants were harvested and, again, IMAC purifications of His_6_-tagged proteins in the supernatant were performed as described above.

Purified protein samples were analysed by xylanase and cellulase activity assays to determine the relative levels of co-purified protein. Reaction rates were calculated by determining the gradient of the linear region of the fluorescence-time plots. The ‘maximum activity’ was set as that detected from co-cultures inoculated with 100% (80 μL) of the strain expressing the C-Xyn-C or C-Cel-C protein and percentage activities were calculated based on this value.

## Supplementary Figures and Tables

**Supplementary Figure 1.**
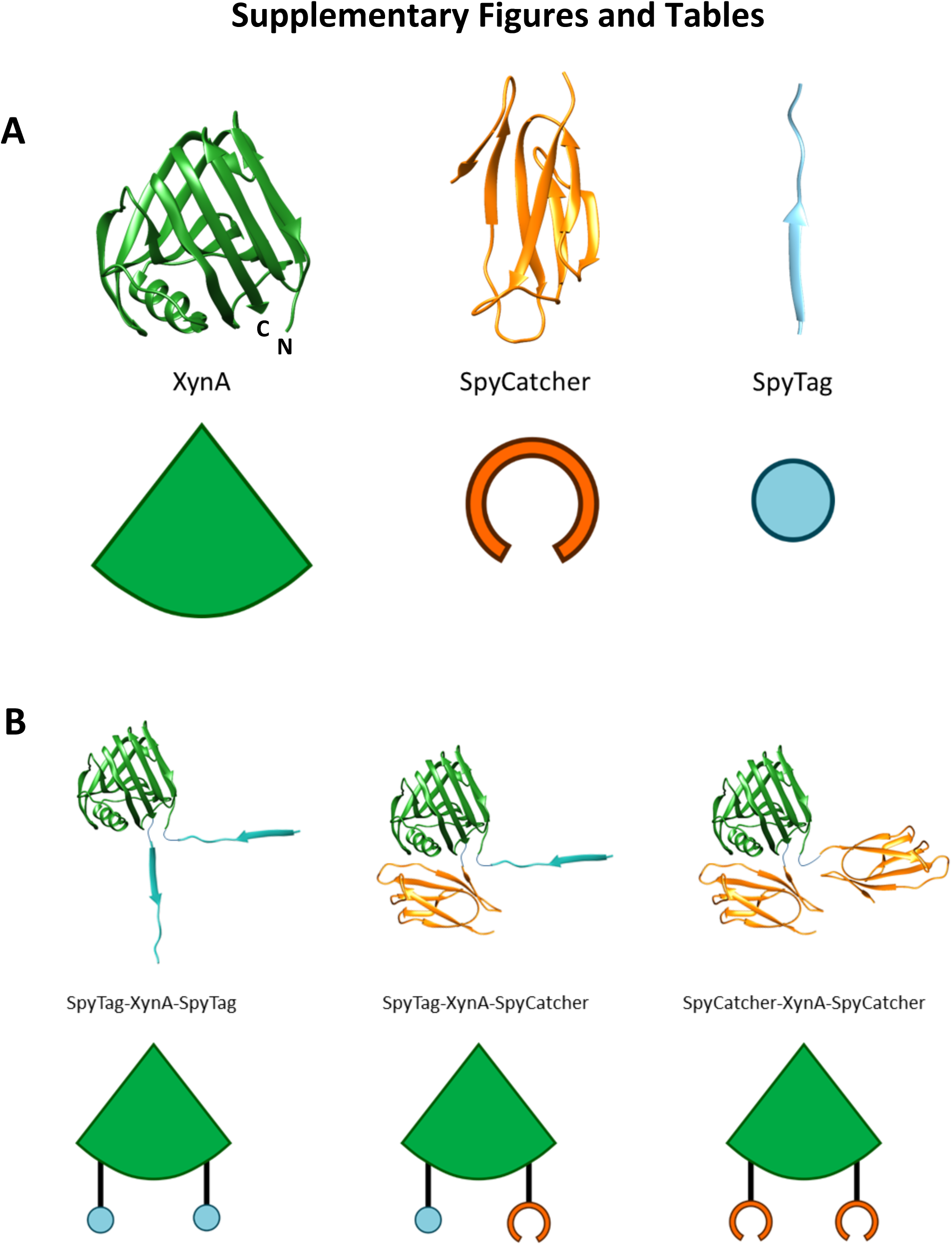
XynA fusion protein schematics. **A** X-ray crystal structures of XynA (PDB ID: 2DCY), SpyCatcher and SpyTag (both base on PDB ID: 4MLI) are shown alongside the graphic representations used elsewhere in this study. The N- and C-termini of XynA (following removal of the N-terminal signal peptide) are very close to one another: 3.8 Å as measured between the N-terminal nitrogen and C-terminal C_α_ atoms (using Chimera, UCSF). **B** Approximate structures of XynA-SpyPart fusion proteins and graphic representations used throughout this study. Note that the structures shown were *not* experimental determined and are purely for illustrative purposes. SpyParts are fused the N- and C-termini of XynA through flexible 2 amino acid glycine-serine linkers. Due the close proximity of the SpyParts, cyclisation reactions are possible.

**Supplementary Figure 2.**
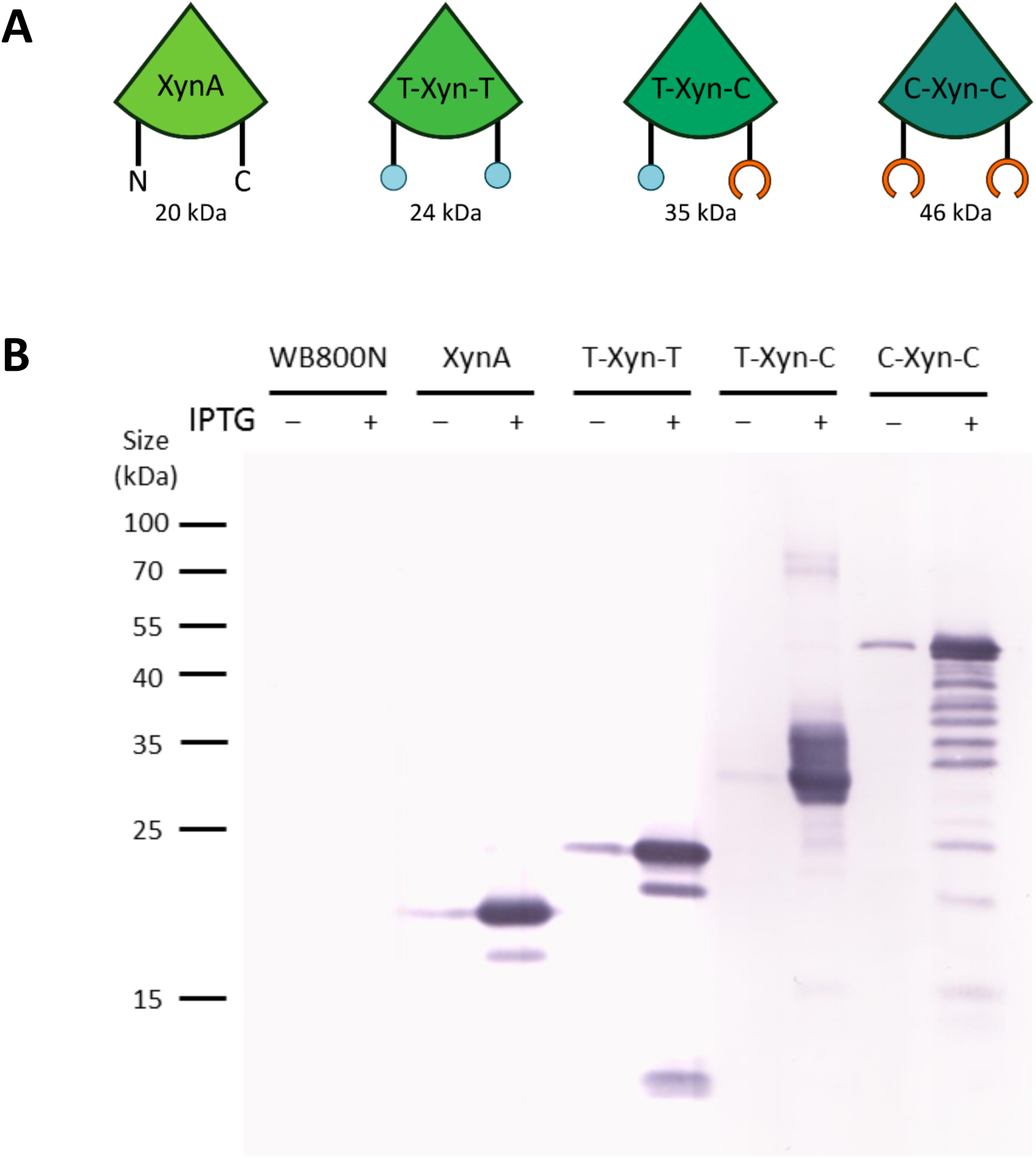
XynA fusion protein construct expression and secretion. **A** Constructs analysed below and their approximate molecular weights. **B** Western blot analysis of secreted XynA fusion proteins. Four fusion proteins based on XynA were designed: XynA, SpyTag-XynA-SpyTag (T-Xyn-T), SpyTag-XynA-SpyCatcher (T-Xyn-C) and SpyCatcher-XynA-SpyCatcher (C-Xyn-C). Each possessed the native XynA signal peptide at the N-terminus as well as a C-terminal His_6_ tag. Each recombinant protein was expressed from the IPTG-inducible *B. subtilis-E. coli* shuttle vector pHT01. After 6 h growth culture supernatants were collected and analysed by SDS-PAGE for protein expression. Supernatants from uninduced (-) and IPTG-induced (+) cultures were 10x concentrated by TCA precipitation prior to Western blotting using an anti-His_6_ primary antibody. The background strain *B. subtilis* WB800N possessing no plasmid is included as a negative control. All constructs are well-expressed and secreted. There is also clearly some leaky expression from the P_grac_ promoter even in the absence of IPTG induction. In addition all constructs are subject to varying degrees of proteolysis – resulting in lower molecular weight species.

**Supplementary Figure 3.**
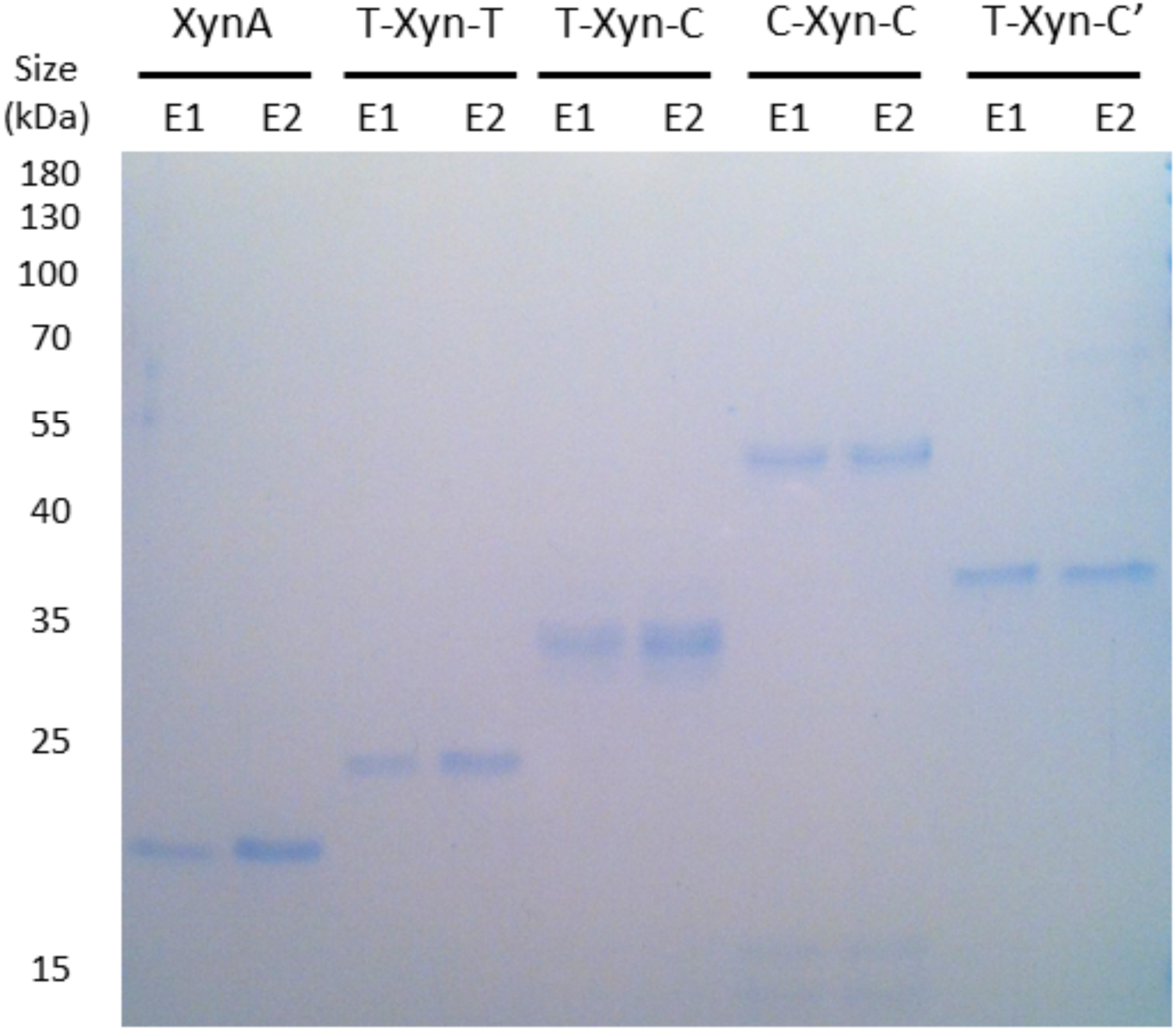
SDS-PAGE analysis of purified XynA fusion proteins. Various XynA fusion proteins were purified from the culture supernatant by IMAC purification: the full-length xylanase XynA, SpyTag-XynA-SpyTag (T-Xyn-T), SpyTag-XynA-SpyCatcher (T-Xyn-C), SpyCatcher-XynA-SpyCatcher (C-Xyn-C) and SpyTag-XynA-SpyCatcher^E77Q^ (T-Xyn-C’). Samples were bound to the Ni-NTA resin and after washing were eluted in two separate 250 μL fractions, E1 and E2. We added 4 μL of 5x SDS-PAGE sample buffer to 16 μL of E1 and E2 samples and then boiled for 10 min. 10 μL samples were loaded and separated by SDS-PAGE with subsequent Coomassie staining. Purification was successful for each protein, with little detectable contaminating species present. Since roughly equal yields were obtained in both the E1 and E2 fractions for each protein, the fractions were pooled for downstream investigations. Approximate molecular weights of each species were: XynA ∼20 kDa, T-Xyn-T ∼ 24 kDa, T-Xyn-C ∼35 kDa, C-Xyn-C ∼46 kDa and T-Xyn-C’ ∼35 kDa. The difference in mobility of T-Xyn-C and T-Xyn-C’ proteins due to cyclisation is very apparent here.

**Supplementary Figure 4.**
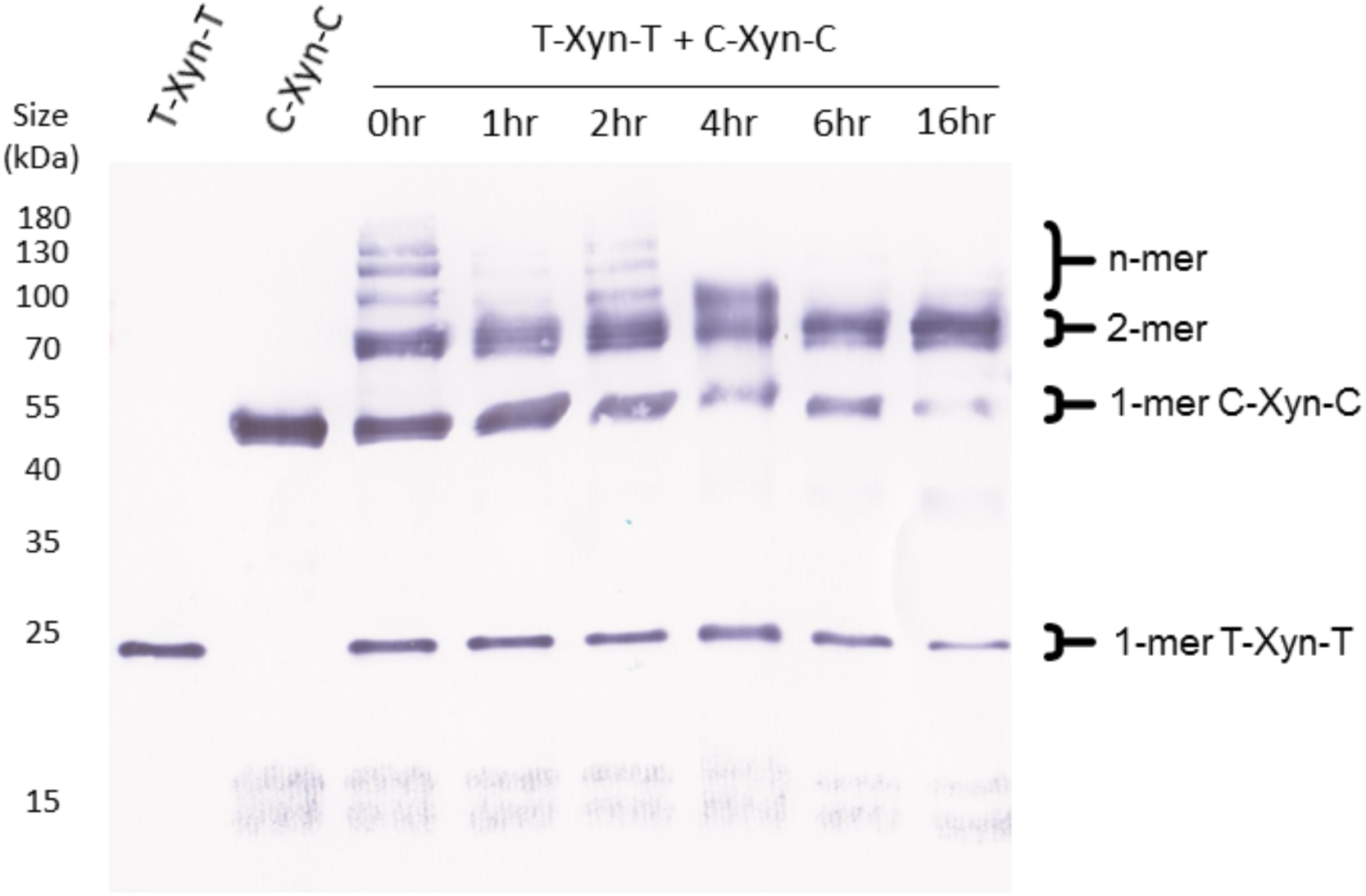
Western blot analysis of purified XynA fusion proteins. SpyTag-XynA-SpyTag (T-Xyn-T) and SpyCatcher-XynA-SpyCatcher (C-Xyn-C) were purified individually from culture supernatant. The T-Xyn-T and C-Xyn-C proteins were mixed 1:1 in a total volume of 20 μL and incubated at 25°C for indicated time periods, after which 5 μL of 5x SDS-PAGE sample buffer was added and samples boiled for 10 min. Western blotting was performed on all samples as well as unmixed samples of each protein. Figure 2B is a cropped version of this blot. Clear evidence of polymerisation (high molecular weight species) is evident even when mixed samples were boiled immediately after mixing. Interestingly, over time, the presence of species larger than dimers appears to decrease, while dimers appear to constantly increase in abundance. The reasons for this are not clear. However, we speculate that, at the point of linkage between two proteins, there is a bifurcation between two pathways: circularisation and linear polymerisation. Since larger, polymerised species tend to be less efficiently transferred to the PVDF membrane during Western blotting, linear polymer species disappear over time as their size grows. Circularised dimers in contrast cannot grow larger over time and so appear to steadily increase in abundance. Regardless, it is clear that over time increasing amounts of monomer species become conjugated, with the reaction reaching near-completion (consumption of all of the C-Xyn-C monomer) after 16 h. Approximate molecular weights of each species were: T-Xyn-T ∼ 24 kDa, C-Xyn-C ∼46 kDa and T-Xyn-T + C-Xyn-C dimers ∼70 kDa.

**Supplementary Figure 5.**
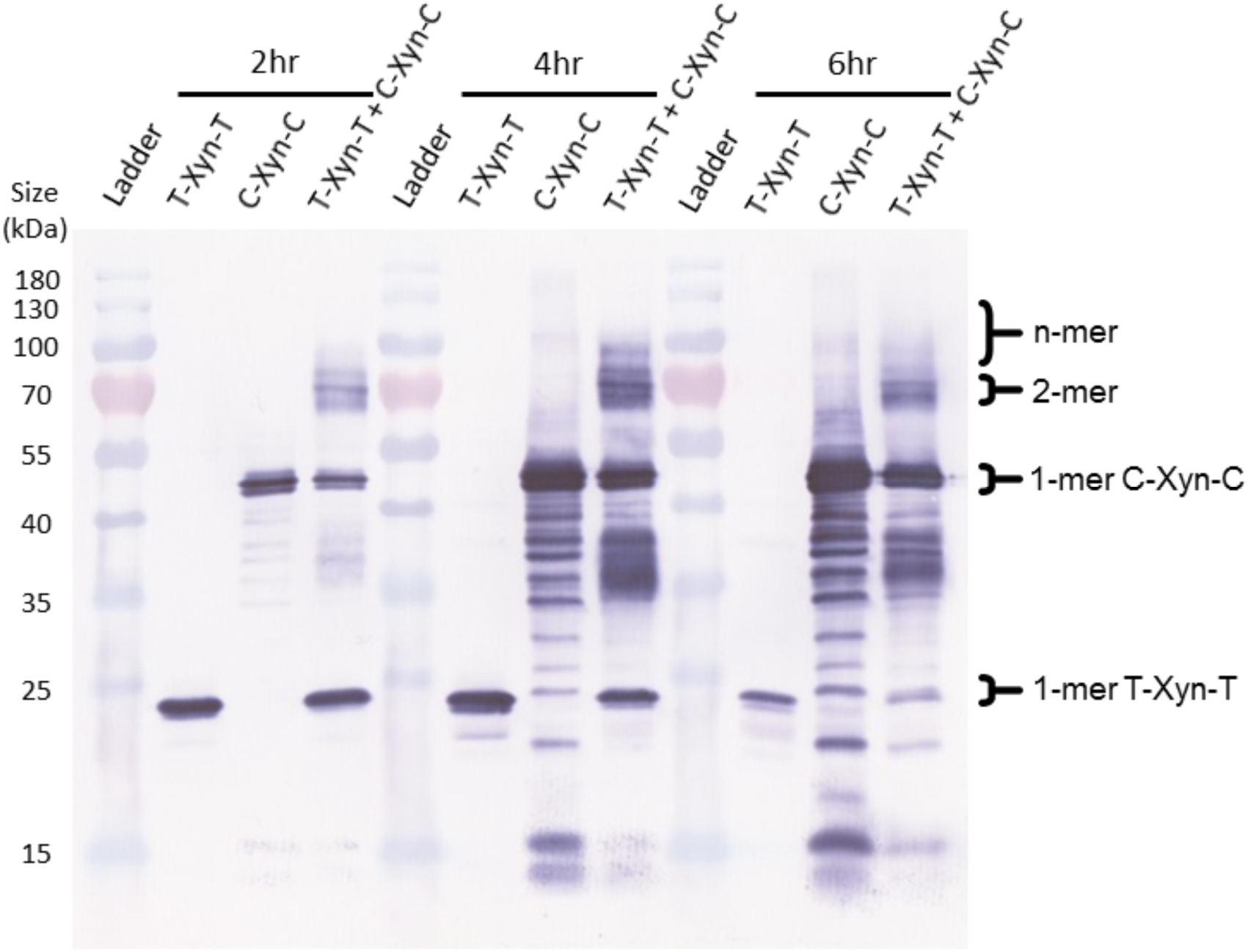
Western blot analysis of co-cultured XynA fusion proteins. Strains expressing SpyTag-XynA-SpyTag (T-Xyn-T) and SpyCatcher-XynA-SpyCatcher (C-Xyn-C) were grown alone and in coculture. At various time points, supernatant samples were harvested, concentrated 10x by TCA precipitation and analysed by Western blotting using an anti-His_6_ antibody. Figure 2C is a cropped version of this blot. Samples were run alongside the PageRuler Prestained Protein Ladder (Thermo) which produces light blue and red bands. After 2 h of coculture, clear evidence of dimeric species can be seen. However, longer incubations produce less clear results due to the high levels of proteolysis, particularly for the CXC construct. At 4 h dimeric and possibly trimeric species are apparent. After 6 h smearing and proteolysis prevents reliable data interpretation. However, the majority of the monomeric T-Xyn-T protein appears to have been consumed after 6 h, indicating that the SpyTag-SpyCatcher conjugation is near-complete. Approximate molecular weights of each species were: T-Xyn-T ∼ 24 kDa, C-Xyn-C ∼46 kDa and T-Xyn-T + C-Xyn-C dimers ∼70 kDa.

**Supplementary Figure 6.**
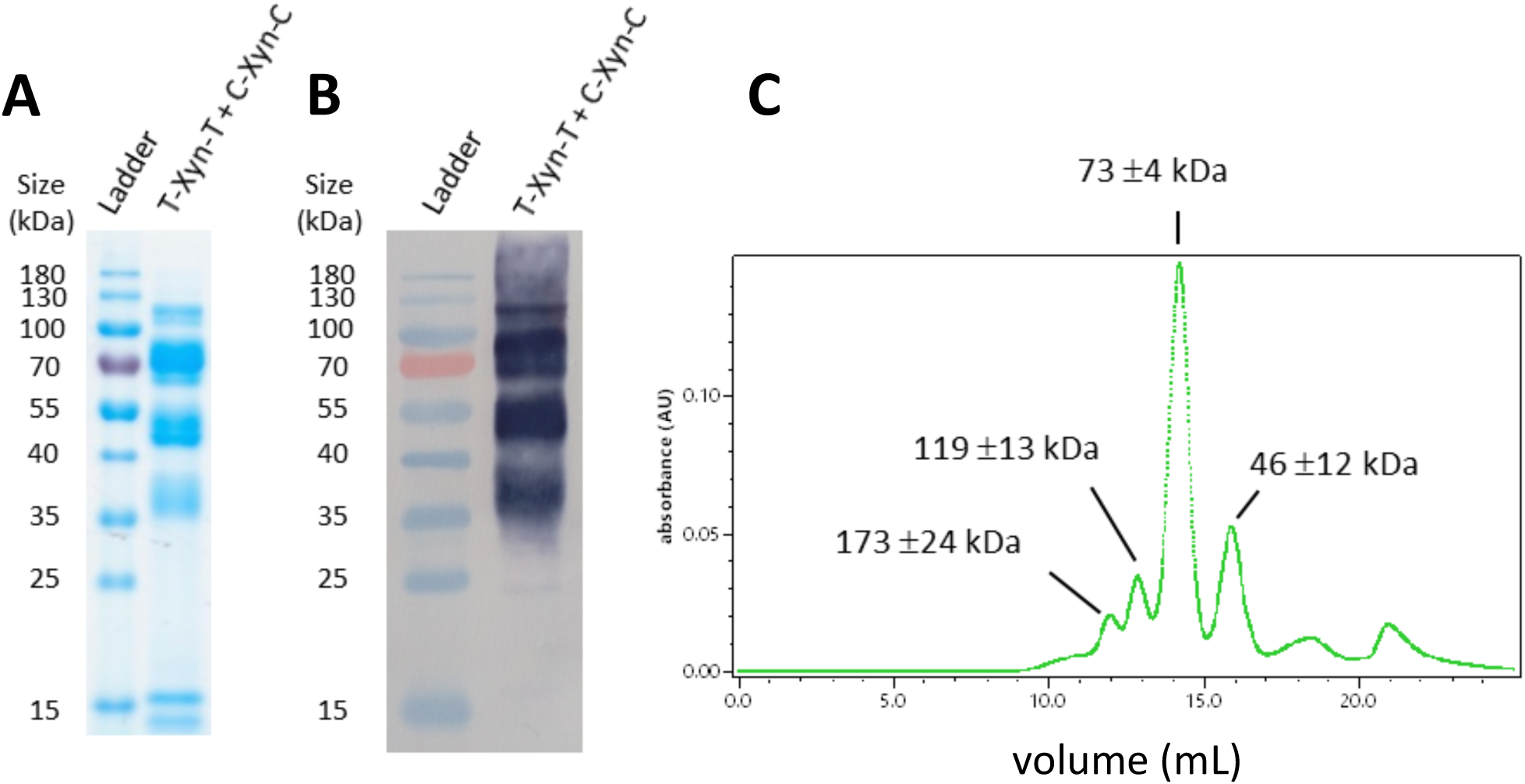
Size exclusion chromatography multi-angle light scattering (SEC-MALS) analysis of purified T-Xyn-T and C-Xyn-C co-culture conjugates. **A** SDS-PAGE analysis with Coomassie staining and **B** Western blotting with an anti- His_6_ antibody indicated that the first elution fraction contained multiple oligomerised forms of T-Xyn-T and C-Xyn-C. In both cases samples were run alongside the PageRuler Prestained Protein Ladder (Thermo) which produces light blue and red bands. Not all species could be reliably identified – for instance, the 37 kDa band lies between the sizes expected for either monomer (T-Xyn-T ∼24 kDa and C-Xyn-C ∼ 46 kDa). Notably, it appears that almost all of the T-Xyn-T protein has reacted. **C** To corroborate these findings, a sample of the first elution fraction was analysed by SEC-MALS instrument (Wyatt) with a flow rate of 0.5 ml.min^−1^ and a 1 sec data collection interval. Data were analysed with Astra 6. Here the UV-vis absorbance trace is shown over various elution volumes annotated with approximate molecular weights calculated based on light scattering data. The major species present has a molecular weight corresponding to a T-Xyn-T + C-Xyn-C dimer (∼70kDa). Additional larger forms were detected, however, molecular weights could not be estimated with sufficient certainty to reliably classify these as specific oligomers.

**Supplementary Figure 7.**
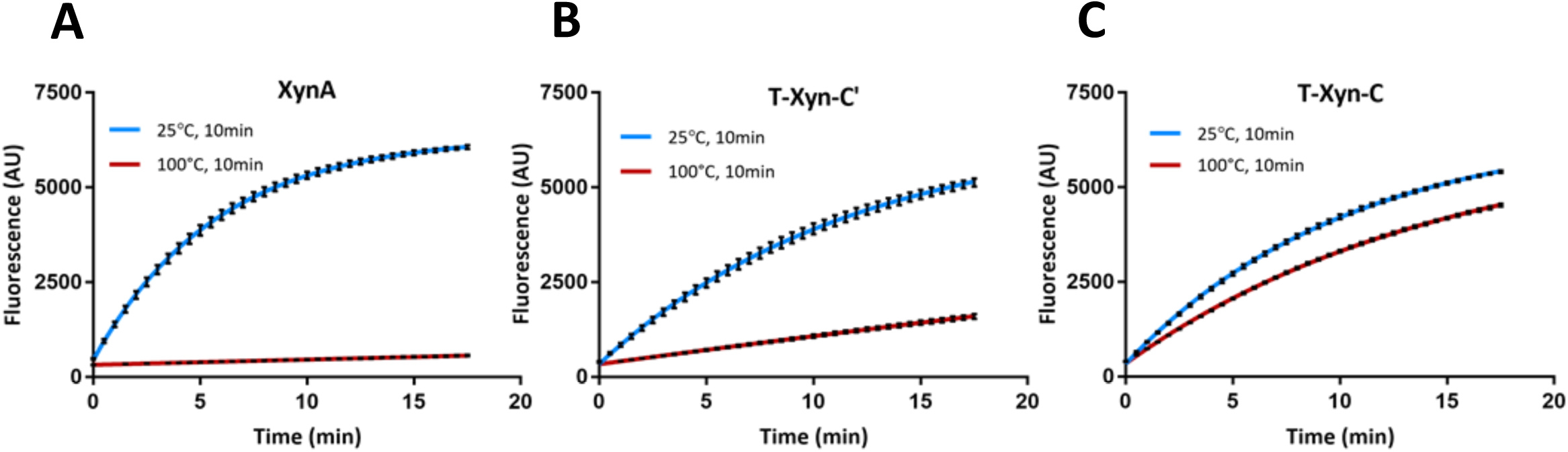
Representative xylanase assay traces following high-temperature exposure. Here culture supernatants from strains expressing **D** XynA, **E** T-Xyn-C’ and **F** T-Xyn-C were subjected to 25°C or 100°C for 10 minutes, cooled to 4°C then equilibrated to room temperature and finally assayed for xylanase activity. Product formation rates were calculated over the linear region of the curves to produce the data in Figure 3F. Samples were analysed in triplicate, data represent the mean ± 1 SD.

**Supplementary Figure 8.**
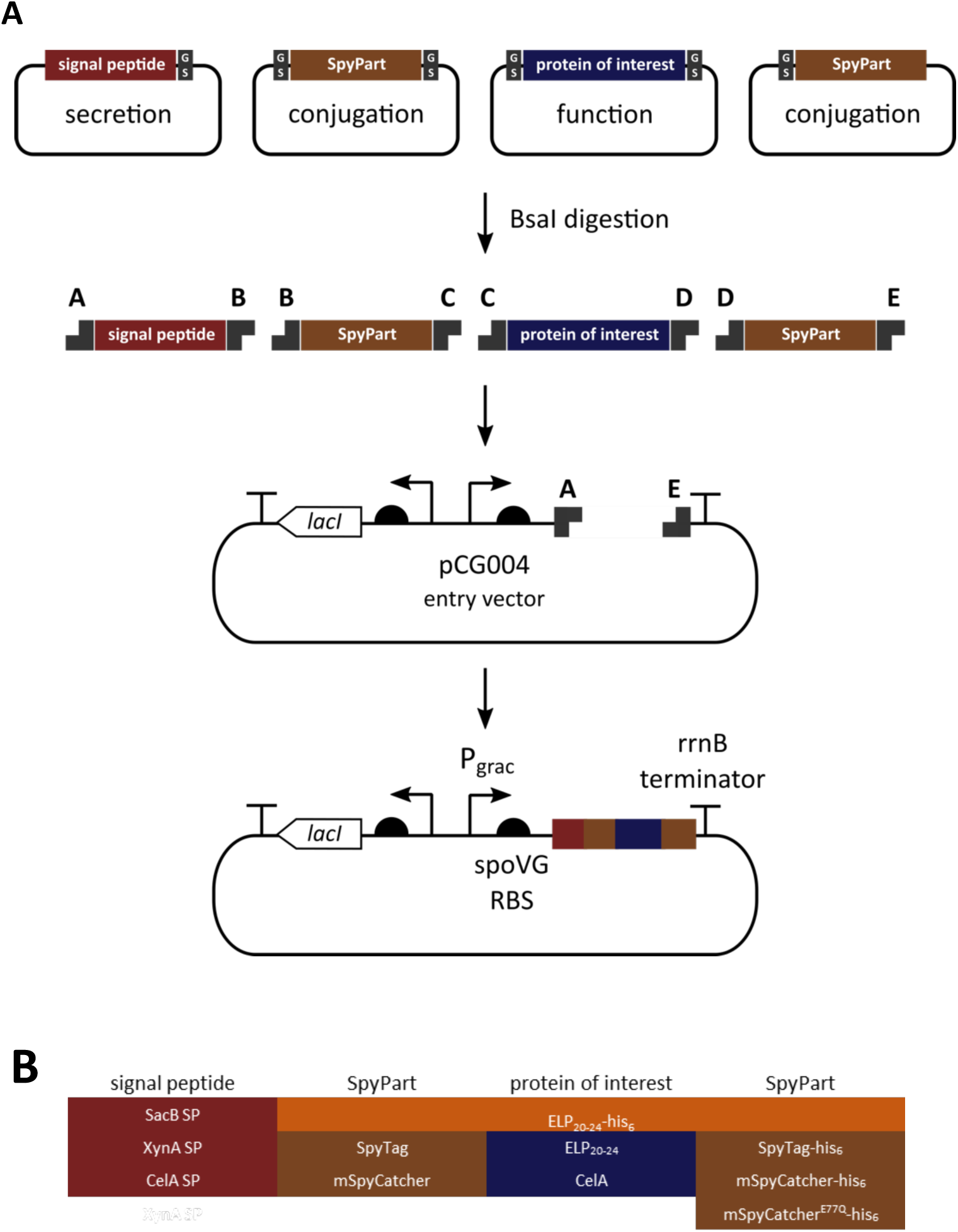
Golden Gate assembly method design and composition. **A** Protein modules were pre-cloned into the part vector pYTK001, sequence-verified and stocked. Four separate positions within the final ORF were defined based on the sequence of upstream and downstream 4 bp overhangs generated by BsaI digestion. Signal peptide parts were cloned with an upstream ‘A’ and downstream ‘B’ overhang, upstream SpyParts were cloned with an upstream ‘B’ and downstream ‘C’ overhang, protein of interest parts were cloned with an upstream ‘C’ and downstream ‘D’ overhang and downstream SpyParts were cloned with an upstream ‘D’ and downstream ‘E’ overhang. Internal overhangs were designed to incorporate two amino acid glycine-serine linkers between ORF parts. Assembly of ORF parts into an entry vector based on pHT01 (pCG004) enabled one-step construction of protein expression constructs. pCG004 possesses a ‘dropout’ part downstream of the P_grac_ promoter and spoVG ribosome binding site (RBS) and upstream of the rrnB terminator. The dropout part consists of a constitutive GFP expression cassette flanked by BsaI restriction sites producing an upstream ‘A’ overhang and downstream ‘E’ overhang. Successful Golden Gate assemblies will therefore result in removal of the GFP expression cassette enabling visual (green-white) screening of transformants. **B** Protein module parts cloned and verified during the course of this study – additional parts not used in this study have also been created. Where required, 4 bp overhangs can be modified to allow assembly of 2 or 1 ORF parts into the entry vector (e.g. the ELP_20-24_-His _6_ part was cloned with an upstream ‘B’ overhang and downstream ‘E’ overhang). Sequences of selected parts and the entry vector pCG004 are included in Supplementary Table 4 and deposited on Addgene.

**Supplementary Figure 9.**
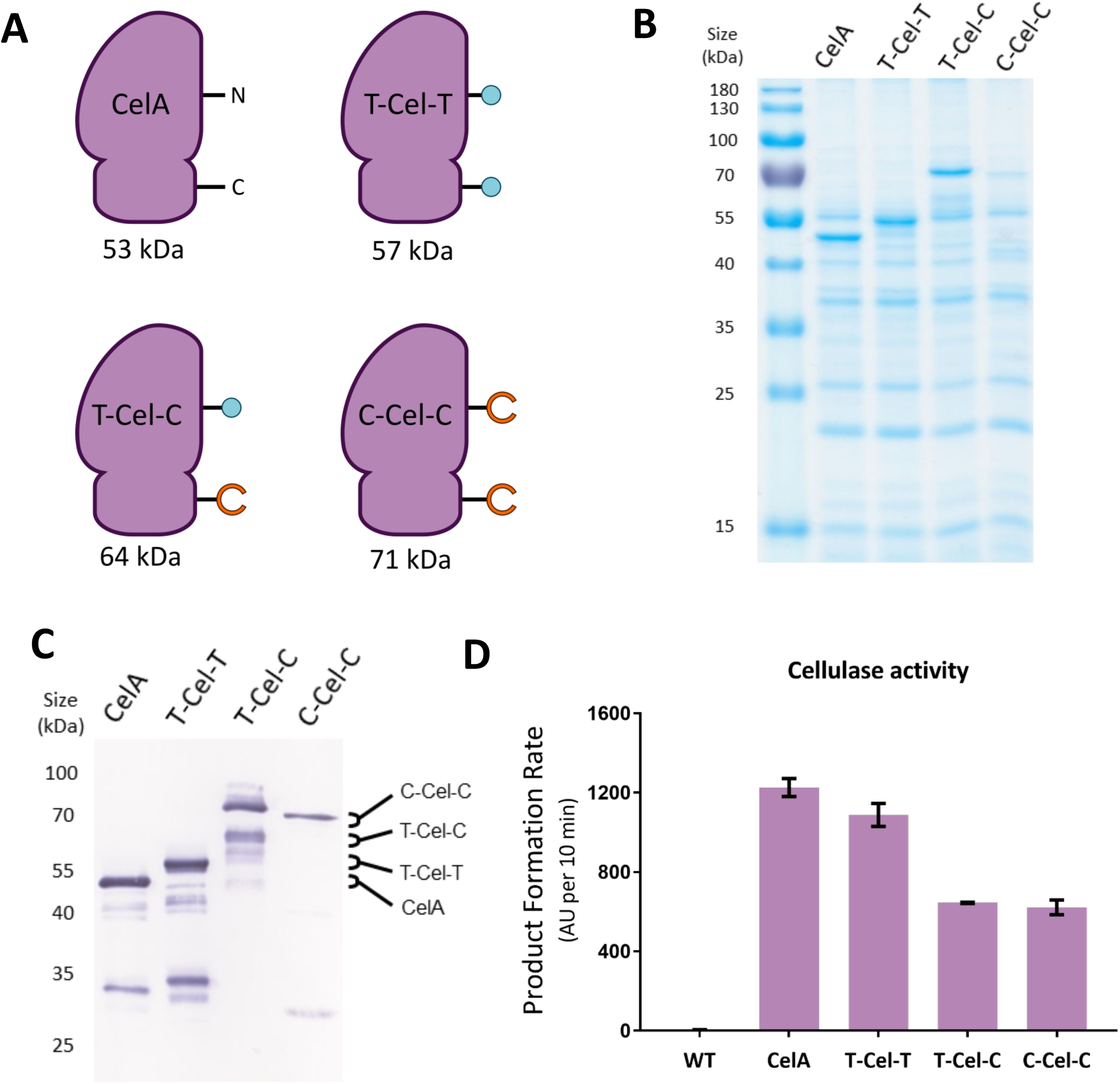
Design, secretion and activity of SpyPart-CelA fusion proteins. **A** Four recombinant proteins based on CelA were designed: the full-length cellulase CelA, SpyTag-CelA-SpyTag (T-Cel-T), SpyTag-CelA-SpyCatcher (T-Cel-C) and SpyCatcher-CelA-SpyCatcher (C-Cel-C). Cartoons were specifically designed to reflect the two-domain architecture of CelA. Each possessed the native CelA signal peptide at the N-terminus as well as a C-terminal His_6_ tag. Each recombinant protein was expressed from the IPTG-inducible *B. subtilis-E. coli* shuttle vector pHT01 and culture supernatant analysed by SDS-PAGE for protein expression. Supernatants from IPTG-induced cultures grown for 6 h were 10x concentrated by TCA precipitation prior to **B** SDS-PAGE with Coomassie staining and **C** western blot analysis. Western blotting was performed using an anti-His_6_ primary antibody. All constructs were well-expressed at roughly equal yields, although some proteolytic degradation is evident. Interestingly, the majority of the T-Cel-C protein appears to have lower than expected mobility. This is consistent with cyclisation through reaction of the SpyTag and SpyCatcher. **D** Culture supernatants were analysed for cellulase activity using a fluorogenic substrate. Product formation rates were calculated over the first minute from triplicate samples, data represent the mean ± 1 SD.

**Supplementary Figure 10.**
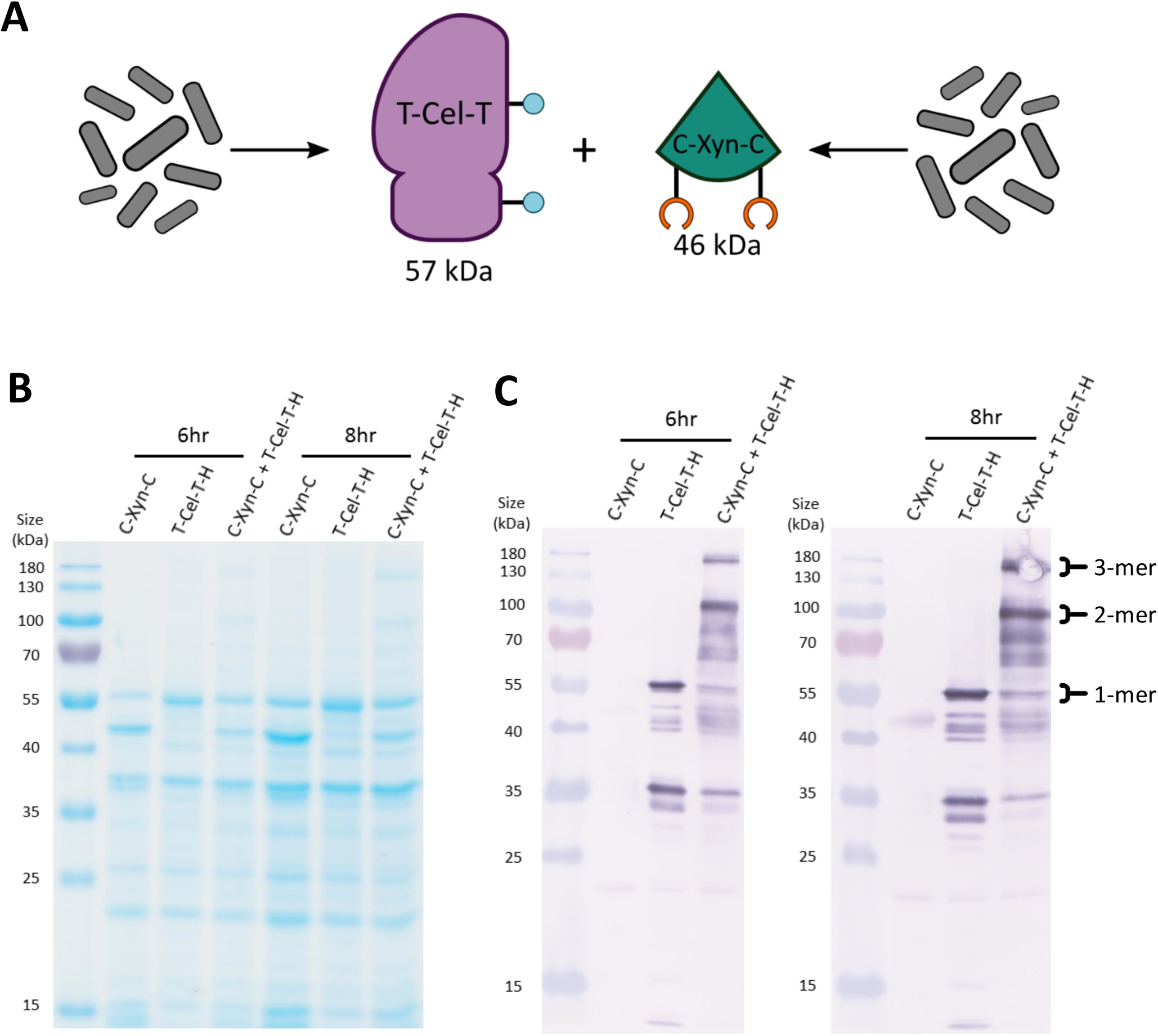
Two-strain co-cultures of T-Cel-T and C-Xyn-C produce multi-protein complexes. **A** Schematic showing the two-strain coculture setup and design of SpyTag-CelA-SpyTag-His_6_ (T-Cel-T) and SpyCatcher-XynA-SpyCatcher (C-Xyn-C). Mono-cultures and a two-strain co-culture of strains expressing each protein were prepared. Supernatant samples were collected after incubation for 6 h and 8 h and concentrated 10x by TCA precipitation. Concentrated samples were analysed by **B** SDS-PAGE with Coomassie staining and **C** Western blotting with an anti- His_6_ primary antibody. Notably, since only the T-Cel-T construct possesses a His_6_ tag, the C-Xyn-C protein is not detectable by Western blotting. While SDS-PAGE analysis produces relatively faint bands, clear evidence of conjugation of T-Cel-T and C-Xyn-C is visible by Western blotting. Species with apparent molecular weights corresponding to dimers and trimers are discernible (expected molecular weights: T-Cel-T + C-Xyn-C ∼103 kDa, T-Cel-T + 2(C-Xyn-C) ∼149 kDa and 2(T-Cel-T) + C-Xyn-C ∼160kDa).

**Supplementary Figure 11.**
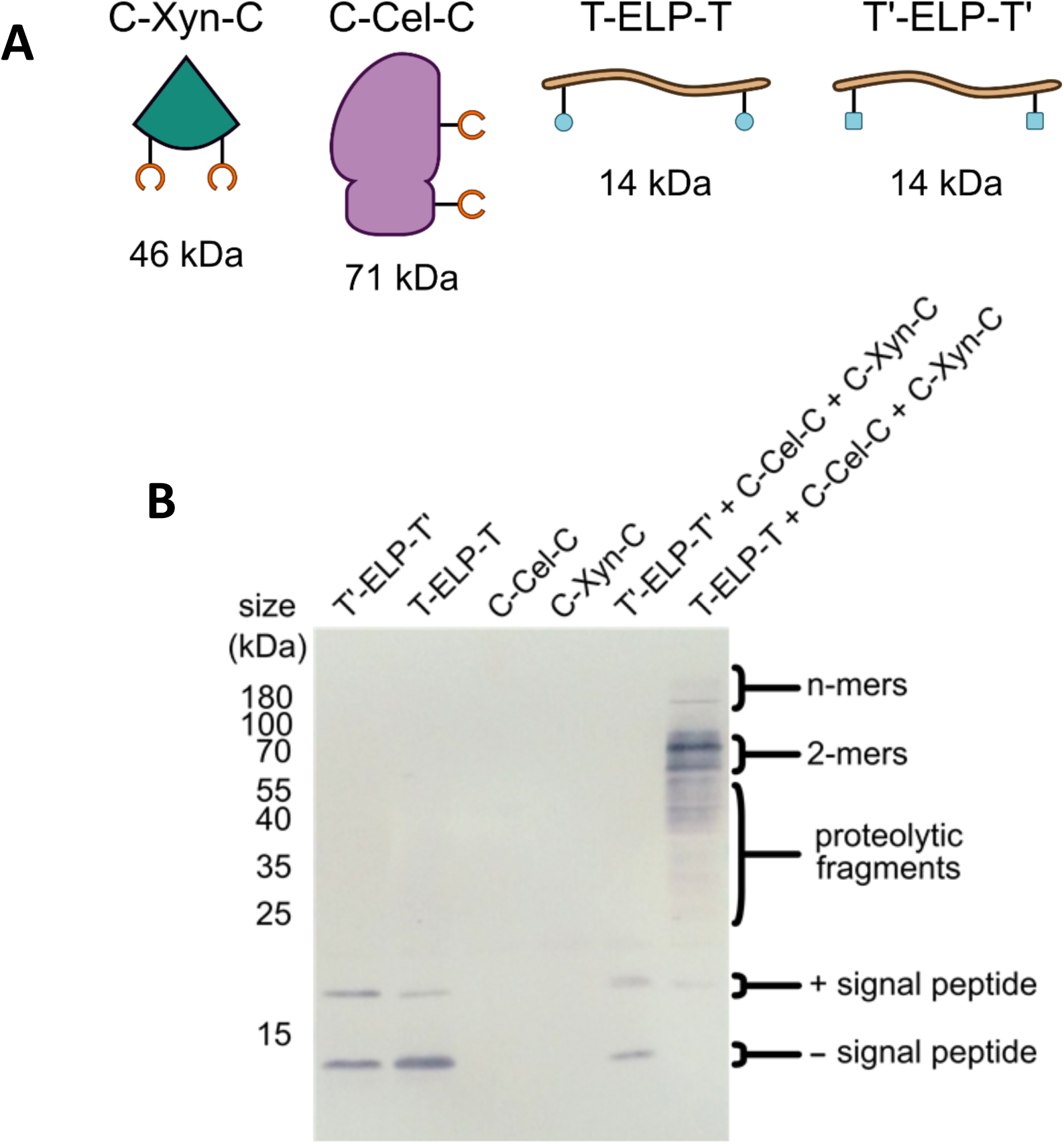
Multi-protein complex formation from three-strain co-cultures. **A** The four recombinant proteins used here: SpyCatcher-XynA-SpyCatcher lacking a C-terminal His_6_ tag (C-Xyn-C), SpyCatcher-CelA-SpyCatcher lacking a C-terminal His_6_ tag (C-Cel-C), SpyTag-ELP_20-24_-SpyTag-His_6_ (T-ELP-T) and SpyTag^DA^-ELP_20-24_-SpyTag^DA^-His_6_ (T’-ELP-T’). Molecular weights of each species are also given (calculated assuming removal of N-terminal signal peptides). **B** Culture supernatants from strains expressing each of the four recombinant proteins were analysed by Western blotting with an anti-His _6_ antibody. Since only the ELP_20 -24_-containing constructs possess C-terminal His_6_ tags, only these are detected. Both T-ELP-T and T’-ELP-T’ well expressed and secreted at similar levels. Under three-strain co-cultures, almost the entirety of T-ELP-T is incorporated into multi-protein complexes with C-Cel-C and C-Xyn-C, an interaction dependent on the SpyTag-SpyCatcher reaction. Proteolysis hampers precise classification of the oligomeric species present.

**Supplementary Table 1.**
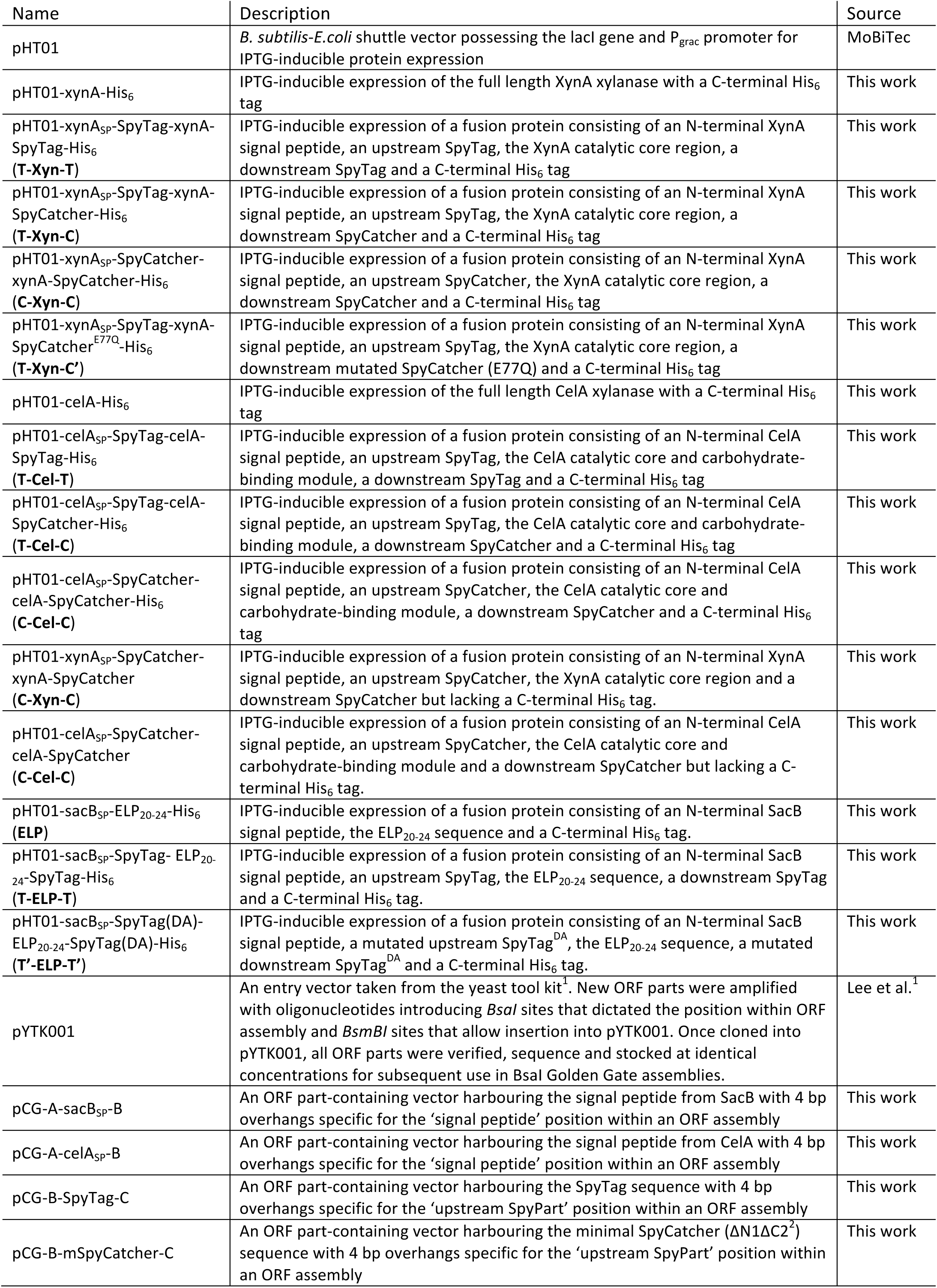
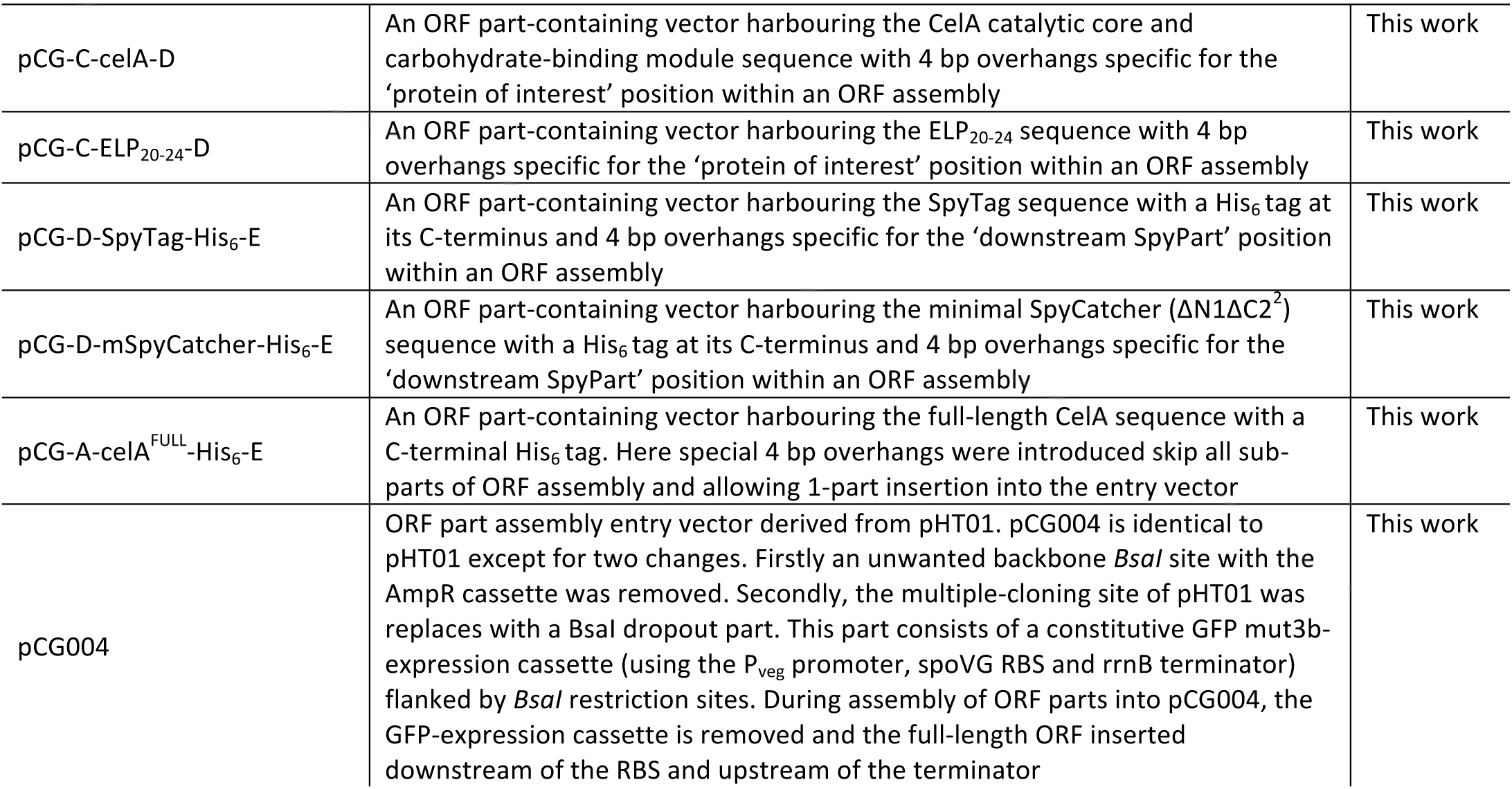
**Plasmids used in this study**

**Supplementary Table 2.** **Strains used in this study**

| Name | Description | Source |
| --- | --- | --- |
| *Bacillus subtilis* WB800N | *trpC2 nprE aprE epr bpr mpr::ble nprB::bsrΔvpr wprA::hyg cm::neo*; NeoR | MoBiTec |
| *Escherichia coli* Turbo | F' *proA<sup>+</sup>B<sup>+</sup> lacI<sup>q</sup> ΔlacZM15 / fhuA2 Δ(lac-proAB) glnV galK16 galE15 R(zgb-210::Tn10)*Tet<sup>S</sup> *endA1 thi-1 Δ(hsdS-mcrB)5* | NEB |

**Supplementary Table 3.**
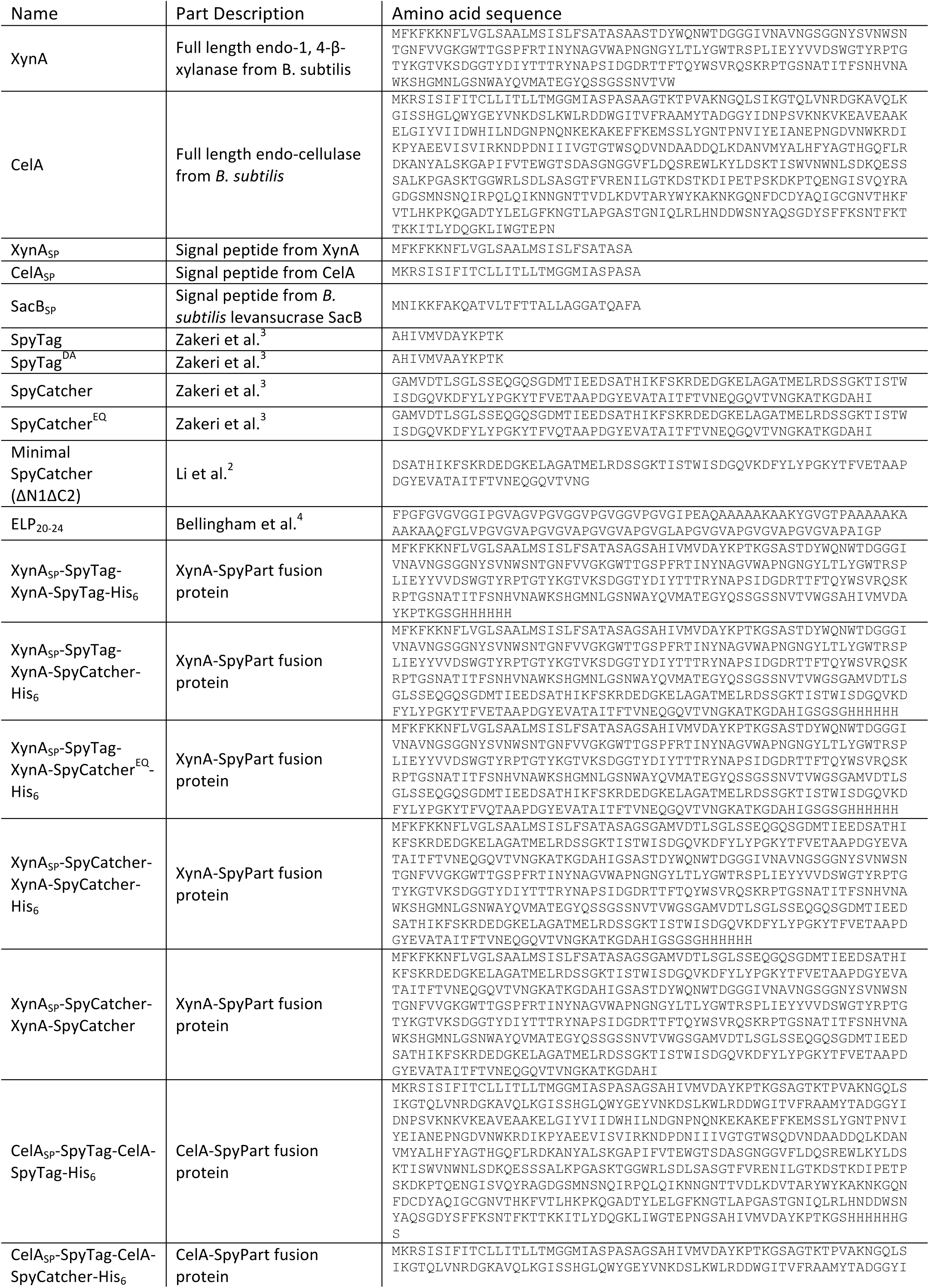
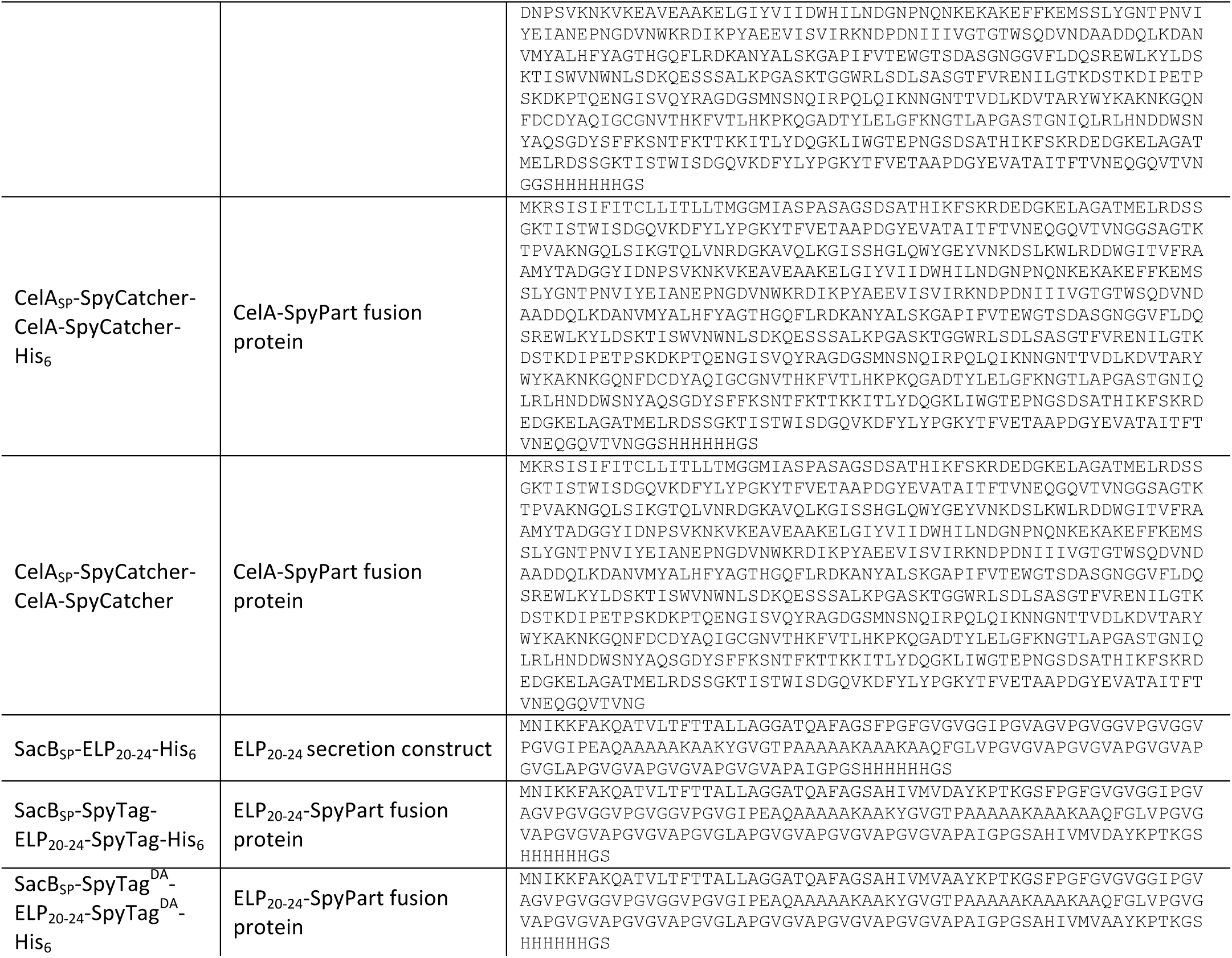
**Amino acid sequences of protein parts used in this study**

**Supplementary Table 4.**
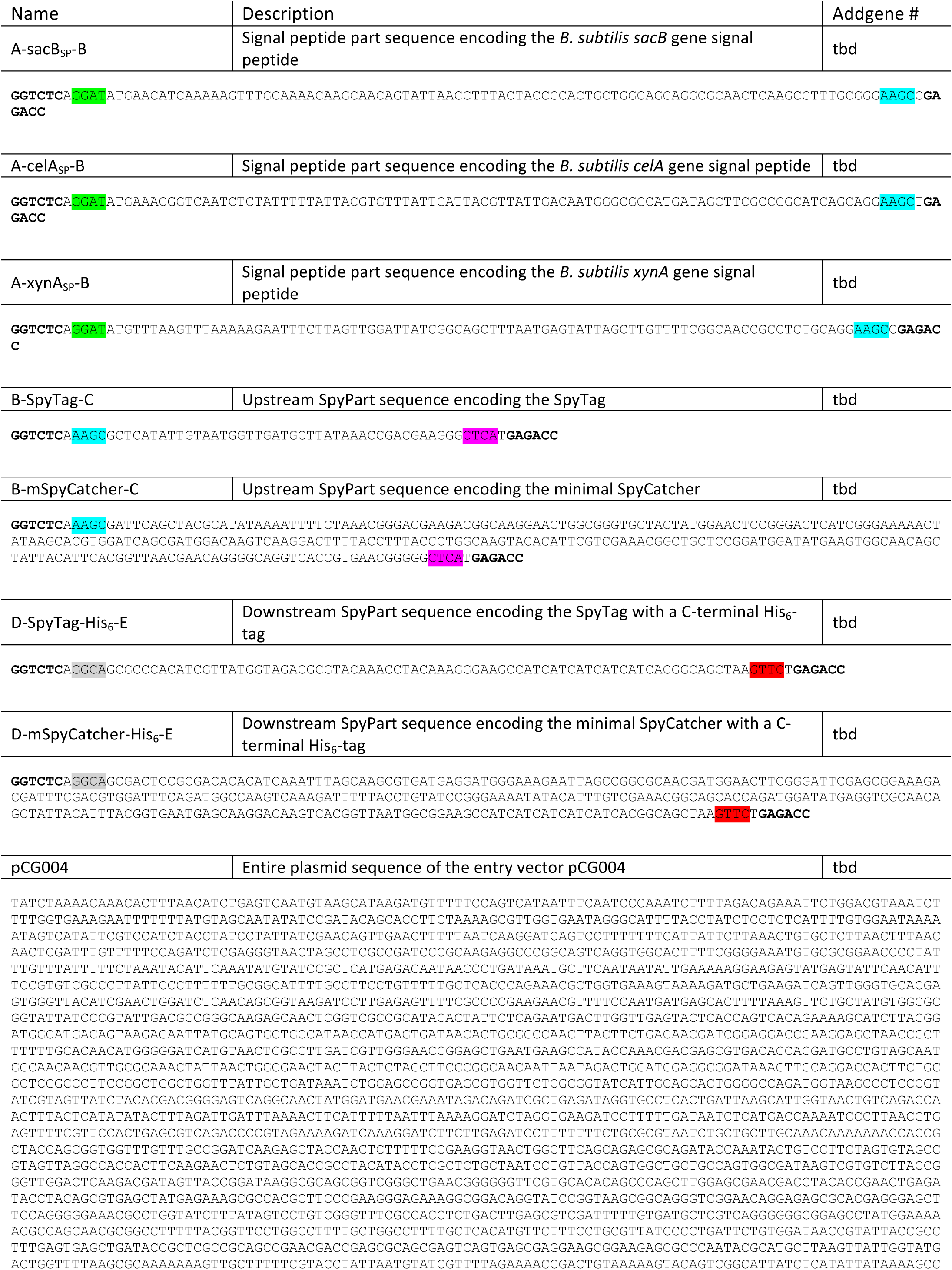
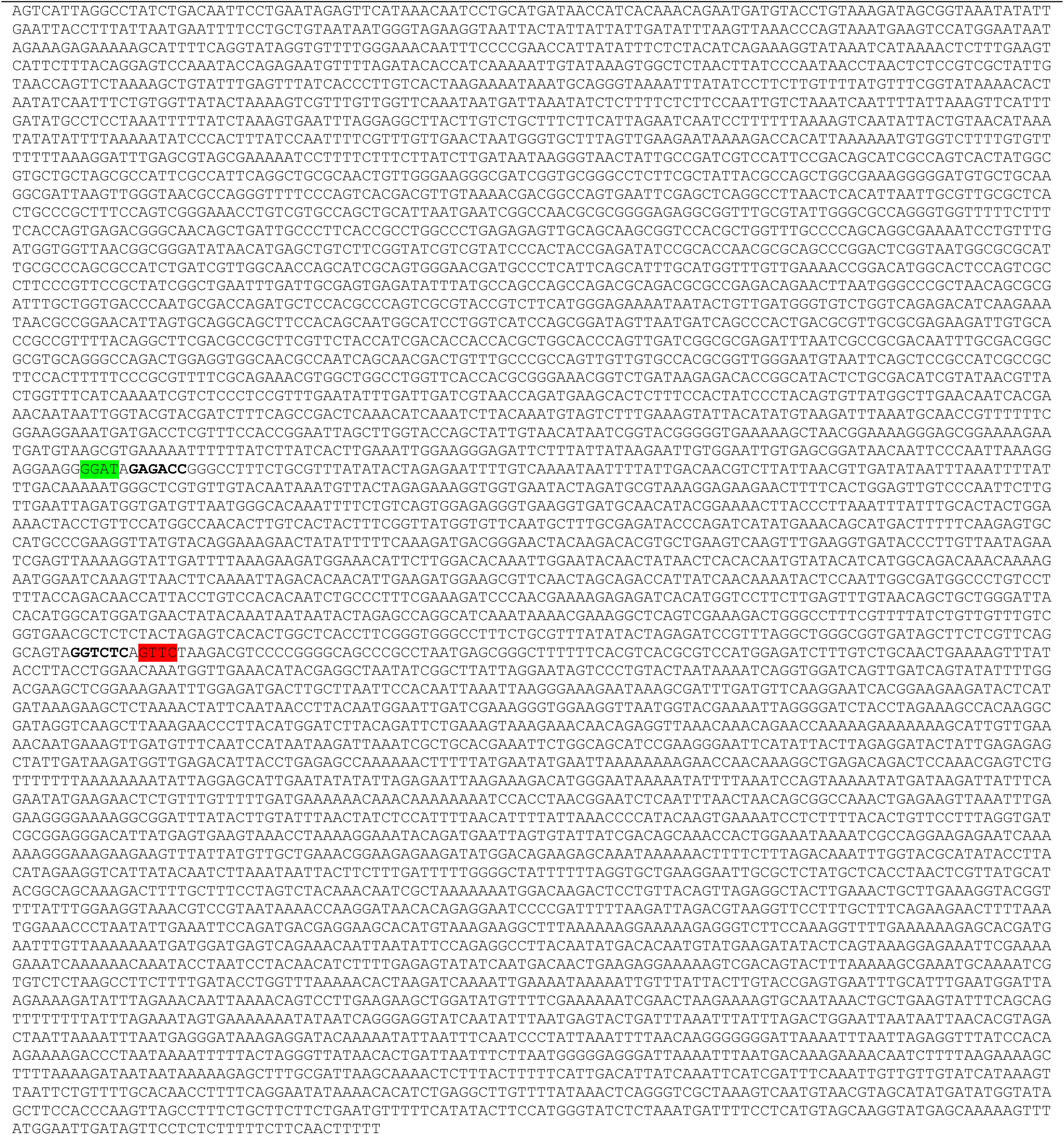
**Golden gate assembly parts used in this study and available from Addgene.** BsaI restriction enzyme recognition sites are shown in bold and overhangs are highlighted.

## REFERENCES

(1) Parmar, P. a., Chow, L. W., St-Pierre, J.-P., Horejs, C.-M., Peng, Y. Y., Werkmeister, J. a., Ramshaw, J. a. M., and Stevens, M. M. (2015) Collagen-mimetic peptide-modifiable hydrogels for articular cartilage regeneration. Biomaterials 54, 213–225.

(2) Minamihata, K., Yamaguchi, S., Nakajima, K., and Nagamune, T. (2016) Tyrosine Coupling Creates a Hyperbranched Multivalent Protein Polymer Using Horseradish Peroxidase via Bipolar Conjugation Points. Bioconjug. Chem. 27, 1348–1359.

(3) Vidal, G., Blanchi, T., Mieszawska, A. J., Calabrese, R., Rossi, C., Vigneron, P., Duval, J. L., Kaplan, D. L., and Egles, C. (2013) Enhanced cellular adhesion on titanium by silk functionalized with titanium binding and RGD peptides. Acta Biomater. 9, 4935–4943.

(4) Chen, A. Y., Deng, Z., Billings, A. N., Seker, U. O. S., Lu, M. Y., Citorik, R. J., Zakeri, B., and Lu, T. K. (2014) Synthesis and patterning of tunable multiscale materials with engineered cells. Nat. Mater. 13, 515–23.

(5) Nguyen, P. Q., Botyanszki, Z., Tay, P. K. R., and Joshi, N. S. (2014) Programmable biofilm-based materials from engineered curli nanofibres. Nat. Commun. 5, 4945.

(6) Myhrvold, C., Polka, J. K., and Silver, P. A. (2016) Synthetic Lipid-Containing Scaffolds Enhance Production by Colocalizing Enzymes. ACS Synth. Biol. acssynbio.6b00141.

(7) Giessen, T. W., and Silver, P. (2016) A catalytic nanoreactor based on in vivo encapsulation of multiple enzymes in an engineered protein nanocompartment. ChemBioChem.

(8) Brune, K. D., Leneghan, D. B., Brian, I. J., Ishizuka, A. S., Bachmann, M. F., Draper, S. J., Biswas, S., and Howarth, M. (2016) Plug-and-Display: decoration of Virus-Like Particles via isopeptide bonds for modular immunization. Sci. Rep. 6, 19234.

(9) Kan, S.-H., Aoyagi-Scharber, M., Le, S. Q., Vincelette, J., Ohmi, K., Bullens, S., Wendt, D. J., Christianson, T. M., Tiger, P. M. N., Brown, J. R., Lawrence, R., Yip, B. K., Holtzinger, J., Bagri, A., Crippen-Harmon, D., Vondrak, K. N., Chen, Z., Hague, C. M., Woloszynek, J. C., Cheung, D. S., Webster, K. A., Adintori, E. G., Lo, M. J., Wong, W., Fitzpatrick, P. A., LeBowitz, J. H., Crawford, B. E., Bunting, S., Dickson, P. I., and Neufeld, E. F. (2014) Delivery of an enzyme-IGFII fusion protein to the mouse brain is therapeutic for mucopolysaccharidosis type IIIB. Proc. Natl. Acad. Sci. U. S. A. 111, 14870–5.

(10) Chen, X., Zaro, J. L., and Shen, W.-C. (2013) Fusion protein linkers: property, design and functionality. Adv. Drug Deliv. Rev. 65, 1357–69.

(11) Yang, H., Liu, L., and Xu, F. (2016) The promises and challenges of fusion constructs in protein biochemistry and enzymology. Appl. Microbiol. Biotechnol.

(12) Albayrak, C., and Swartz, J. R. (2014) Direct polymerization of proteins. ACS Synth. Biol. 3, 353–62.

(13) Zakeri, B., Fierer, J. O., Celik, E., Chittock, E. C., Schwarz-Linek, U., Moy, V. T., and Howarth, M. (2012) Peptide tag forming a rapid covalent bond to a protein, through engineering a bacterial adhesin. Proc. Natl. Acad. Sci.

(14) Zhang, W.-B., Sun, F., Tirrell, D. a, and Arnold, F. H. (2013) Controlling macromolecular topology with genetically encoded SpyTag-SpyCatcher chemistry. J. Am. Chem. Soc. 135, 13988–97.

(15) Veggiani, G., Nakamura, T., Brenner, M. D., Gayet, R. V, Yan, J., Robinson, C. V, and Howarth, M. (2016) Programmable polyproteams built using twin peptide superglues. Proc. Natl. Acad. Sci. U. S. A. 113, 1202–7.

(16) Sun, F., Zhang, W.-B., Mahdavi, A., Arnold, F. H., and Tirrell, D. a. (2014) Synthesis of bioactive protein hydrogels by genetically encoded SpyTag-SpyCatcher chemistry. Proc. Natl. Acad. Sci. U. S. A. 111.

(17) Botyanszki, Z., Tay, P. K. R., Nguyen, P. Q., Nussbaumer, M. G., and Joshi, N. S. (2015) Engineered catalytic biofilms: Site-specific enzyme immobilization onto E. coli curli nanofibers. Biotechnol. Bioeng. 110, 2016–2024.

(18) Gao, X., Fang, J., Xue, B., Fu, L., and Li, H. (2016) Engineering Protein Hydrogels Using SpyCatcher-SpyTag Chemistry. Biomacromolecules acs.biomac.6b00566.

(19) Schoene, C., Fierer, J. O., Bennett, S. P., and Howarth, M. (2014) SpyTag/SpyCatcher cyclization confers resilience to boiling on a mesophilic enzyme. Angew. Chem. Int. Ed. Engl. 53, 6101–4.

(20) Wang, J., Wang, Y., Wang, X., Zhang, D., Wu, S., and Zhang, G. (2016) Enhanced thermal stability of lichenase from Bacillus subtilis 168 by SpyTag/SpyCatcher-mediated spontaneous cyclization. Biotechnol. Biofuels 9, 79.

(21) Schoene, C., Bennett, S. P., and Howarth, M. (2016) SpyRing interrogation : analyzing how enzyme resilience can be achieved with phytase and distinct cyclization chemistries. Sci. Rep. 1–12.

(22) Alves, N. J., Turner, K. B., Daniele, M. A., Oh, E., Medintz, I. L., and Walper, S. A. (2015) Bacterial Nanobioreactors-Directing Enzyme Packaging into Bacterial Outer Membrane Vesicles. ACS Appl. Mater. Interfaces 7, 24963–24972.

(23) Alves, N. J., Turner, K. B., Medintz, I. L., and Walper, S. A. (2016) Protecting enzymatic function through directed packaging into bacterial outer membrane vesicles. Sci. Rep. 6, 24866.

(24) Fierer, J. O., Veggiani, G., and Howarth, M. (2014) SpyLigase peptide-peptide ligation polymerizes affibodies to enhance magnetic cancer cell capture. Proc. Natl. Acad. Sci. U. S. A. 111, E1176–81.

(25) Lakshmanan, A., Farhadi, A., Nety, S. P., Lee-Gosselin, A., Bourdeau, R. W., Maresca, D., and Shapiro, M. G. (2016) Molecular Engineering of Acoustic Protein Nanostructures. ACS Nano 10, 7314–7322.

(26) Mergulhão, F. J. M., Summers, D. K., and Monteiro, G. a. (2005) Recombinant protein secretion in Escherichia coli. Biotechnol. Adv. 23, 177–202.

(27) Azam, A., Li, C., Metcalf, K. J., and Tullman-Ercek, D. (2015) Type III Secretion as a Generalizable Strategy for the Production of Full-Length Biopolymer-Forming Proteins. Biotechnol. Bioeng. 9999, 1–8.

(28) Zhou, K., Qiao, K., Edgar, S., and Stephanopoulos, G. (2015) Distributing a metabolic pathway among a microbial consortium enhances production of natural products. Nat. Biotechnol. 33, 377–383.

(29) Hays, S. G., Patrick, W. G., Ziesack, M., Oxman, N., and Silver, P. A. (2015) Better together: Engineering and application of microbial symbioses. Curr. Opin. Biotechnol. 36, 40–49.

(30) Brenner, K., You, L., and Arnold, F. H. (2008) Engineering microbial consortia: a new frontier in synthetic biology. Trends Biotechnol. 26, 483–9.

(31) Jones, J. A., Vernacchio, V. R., Sinkoe, A. L., Collins, S. M., Ibrahim, M. H. A., Lachance, D. M., Hahn, J., and Koffas, M. A. G. (2016) Experimental and computational optimization of an Escherichia coli co-culture for the efficient production of flavonoids. Metab. Eng. 35, 55–63.

(32) Harwood, C. R., and Cranenburgh, R. (2008) Bacillus protein secretion: an unfolding story. Trends Microbiol. 16, 73–79.

(33) van Dijl, J. M., and Hecker, M. (2013) Bacillus subtilis: from soil bacterium to super-secreting cell factory. Microb. Cell Fact. 12, 3.

(34) Scott, C. P., Abel-Santos, E., Wall, M., Wahnon, D. C., and Benkovic, S. J. (1999) Production of cyclic peptides and proteins in vivo. Proc. Natl. Acad. Sci. 96, 13638–13643.

(35) Trabi, M., and Craik, D. J. (2002) Circular proteins - no end in sight. Trends Biochem. Sci. 27, 132–138.

(36) Iwai, H., and Plu, A. (1999) Circular L-lactamase : stability enhancement by cyclizing the backbone. FEBS Lett. 459, 166–172.

(37) Schoene, C., Bennett, S. P., and Howarth, M. (2016) SpyRings Declassified: A Blueprint for Using Isopeptide-Mediated Cyclization to Enhance Enzyme Thermal Resilience. Pept. Protein Enzym. Des. 1st ed. Elsevier Inc.

(38) Kuhad, R. C., Gupta, R., and Singh, A. (2011) Microbial cellulases and their industrial applications. Enzyme Res. 2011, 280696.

(39) Zhang, X. Z., and Zhang, Y. H. P. (2010) One-step production of biocommodities from lignocellulosic biomass by recombinant cellulolytic Bacillus subtilis: Opportunities and challenges. Eng. Life Sci. 10, 398–406.

(40) Moraïs, S., Barak, Y., Caspi, J., and Hadar, Y. (2010) Cellulase-xylanase synergy in designer cellulosomes for enhanced degradation of a complex cellulosic substrate. MBio 1, 3–10.

(41) Bellingham, C. M., Woodhouse, K. A., Robson, P., Rothstein, S. J., and Keeley, F. W. (2001) Self-aggregation characteristics of recombinantly expressed human elastin polypeptides. Biochim. Biophys. Acta - Protein Struct. Mol. Enzymol. 1550, 6–19.

(42) Kumar, V., Marín-Navarro, J., and Shukla, P. (2016) Thermostable microbial xylanases for pulp and paper industries: trends, applications and further perspectives. World J. Microbiol. Biotechnol. 32, 1–10.

(43) Juturu, V., and Wu, J. C. (2012) Microbial xylanases: Engineering, production and industrial applications. Biotechnol. Adv. 30, 1219–1227.

(44) Fontes, C. M. G. A., and Gilbert, H. J. (2010) Cellulosomes: Highly Efficient Nanomachines Designed to Deconstruct Plant Cell Wall Complex Carbohydrates. Annu. Rev. Biochem. 79, 655–681.

(45) Arai, T., Matsuoka, S., Cho, H.-Y., Yukawa, H., Inui, M., Wong, S.-L., and Doi, R. H. (2007) Synthesis of Clostridium cellulovorans minicellulosomes by intercellular complementation. Proc. Natl. Acad. Sci. U. S. A. 104, 1456–1460.

(46) Moraïs, S., Shterzer, N., Lamed, R., Bayer, E. A., and Mizrahi, I. (2014) A combined cell-consortium approach for lignocellulose degradation by specialized Lactobacillus plantarum cells. Biotechnol. Biofuels 7, 112.

(47) Stahl, S. W., Nash, M. A., Fried, D. B., Slutzki, M., Barak, Y., Bayer, E. A., and Gaub, H. E. (2012) Single-molecule dissection of the high-affinity cohesin-dockerin complex. Proc. Natl. Acad. Sci. 109, 20431–20436.

(48) Scott, S. R., and Hasty, J. (2016) Quorum Sensing Communication Modules for Microbial Consortia. ACS Synth. Biol.

(49) Abe, H., Wakabayashi, R., Yonemura, H., Yamada, S., Goto, M., and Kamiya, N. (2013) Split Spy0128 as a potent scaffold for protein cross-linking and immobilization. Bioconjug. Chem. 24, 242–250.

(50) Woolfson, D. N. (2005) The design of coiled-coil structures and assemblies. Adv. Protein Chem.

(51) Lee, M. E., DeLoache, W. C., Cervantes, B., and Dueber, J. E. (2015) A Highly-characterized Yeast Toolkit for Modular, Multi-part Assembly. ACS Synth. Biol. 150414151809002.

(52) Cormack, B. P., Valdivia, R. H., and Falkow, S. (1996) FACS-optimized mutants of the green fluorescent protein (GFP). Gene 173, 33–38.

(53) Li, L., Fierer, J. O., Rapoport, T. a, and Howarth, M. (2014) Structural analysis and optimization of the covalent association between SpyCatcher and a peptide Tag. J. Mol. Biol. 426, 309–17.

## Supplementary References

1. Lee, M. E., DeLoache, W. C., Cervantes, B. & Dueber, J. E. A Highly-characterized Yeast Toolkit for Modular, Multi-part Assembly. ACS Synth. Biol. 4, (2015).

2. Li, L., Fierer, J. O., Rapoport, T. a & Howarth, M. Structural analysis and optimization of the covalent association between SpyCatcher and a peptide Tag. J. Mol. Biol. 426, 309–17 (2014).

3. Zakeri, B. et al. Peptide tag forming a rapid covalent bond to a protein, through engineering a bacterial adhesin. Proceedings of the National Academy of Sciences 109, E690–E697 (2012).

4. Bellingham, C. M., Woodhouse, K. A., Robson, P., Rothstein, S. J. & Keeley, F. W. Self-aggregation characteristics of recombinantly expressed human elastin polypeptides. Biochim. Biophys. Acta - Protein Struct. Mol. Enzymol. 1550, 6–19 (2001).

